# A Human Biomimetic Intestinal Mucosa Model to Study Gastrointestinal Development and Disease

**DOI:** 10.1101/2024.08.20.608742

**Authors:** Alessandro Dei, Carlemi Calitz, Joep Korsten, Nina Johannesson, Eline Freeze, Allen Eaves, Sharon Louis, Ryan K. Conder, Wing Chang, Dasja Pajkrt, Katja C. Wolthers, Adithya Sridhar, Salvatore Simmini

**Author notes:** These authors contributed equally (Alessandro Dei, Carlemi Calitz, Adithya Sridhar, Salvatore Simmini).

## Abstract

The intestinal mucosa plays a vital role in nutrient absorption, drug metabolism, and pathogen defence. Advances in single-cell technologies have highlighted the specialised roles of various cell types that execute these diverse functions. Aside from intestinal epithelial cells, fibroblasts play an essential role in regulating the extracellular matrix and controlling pro-inflammatory signalling, and antigen-presenting cells (macrophages and dendritic cells) maintain intestinal homeostasis and immune responses. The incorporation of such cellular complexity within the existing *in vitro* models of the human intestine is currently challenging. To address this, we developed a human intestinal model that accurately mimics the mucosal cellular environment comprising intestinal epithelial cells, intestinal fibroblasts, and antigen presenting cells. This model includes co-cultures of adult and foetal cells, facilitating studies on barrier function, inflammation, and viral infections. It replicates extracellular matrix deposition, Paneth cell differentiation, immune interactions, and can be used to model host-pathogen interactions. Our advanced co-culture model improves the physiological relevance of *in vitro* studies, enabling the exploration of epithelial-mesenchymal-immune crosstalk and its role in intestinal health and disease.

## 1. Introduction

The mucosal lining of the human gastrointestinal (GI) tract is the largest mucosal tissue exposed to the external environment. It plays a crucial role in nutrient absorption and the first-pass metabolism of drugs while protecting non-mucosal sites from invading pathogens [1, 2]. The development and maintenance of the GI tract depend not only on the intestinal epithelial cell (IEC) lining but also on intricate interactions between IECs and cells from different germ layers. Intestinal fibroblasts (IFs) form and regulate the dynamic extracellular matrix (ECM) of the intestinal mucosa, providing essential structural support to the IECs [3]. Additionally, IFs secrete immunomodulatory cytokines, which are fundamental not only during development to preserve immune cell tolerance, but also in the establishment of inflammatory response during injury and infections [4]. Furthermore, the intestinal mucosa is the largest immune organ and is a reservoir to 75% of the immune cells in the body. In particular, APCs, macrophages (MΦ) and dendritic cells (DCs), play critical roles in mucosal immunity. The phenotypical and functional plasticity of these cells is central to maintaining intestinal homeostasis and responding to environmental challenges [5, 6]. Under steady-state conditions, MΦ maintain gut health by recognizing and eliminating pathogens that breach the intestinal barrier, clearing apoptotic cells and debris, enabling tolerance to commensal microbiota and food antigens, and preventing unnecessary inflammation through the production of anti-inflammatory cytokines [7]. Similarly, DCs, specialised in sampling luminal antigens, exhibit a tolerogenic phenotype that facilitates immune tolerance and induction of regulatory T cells [8]. However, tissue damage, infections, autoimmune reactions, and certain environmental factors can disrupt this tolerogenic environment, leading to a heightened inflammatory state. During this phase, APCs shift to a pro-inflammatory state, enhancing their antigen-presenting capacity and stimulating the recruitment and activation of effector T cells, promoting robust immune responses to pathogens or harmful antigens in the gut [9].

Human *in vitro* models that include these complex multicellular diversity and interactions are needed for a better understanding of intestinal development and disease states, including inflammation and infection [10]. Although reductionist, such *in vitro* models enable the study of complex molecular mechanisms and high-throughput drug development [11]. Currently, stem cell-derived human intestinal enteroids and organoids are the gold standard *in vitro* models of the human intestine [12, 13]. However, these organoids do not capture the cellular complexity of the intestinal mucosa, as they primarily contain IECs [14]. Moreover, a significant concern with these three-dimensional (3D) models is their short lifespan upon differentiation, ranging from less than 72 hours to 7 days, due to the terminal differentiation of IECs [1]. Furthermore, these 3D models have a closed cystic orientation that limits their utility for applications that focus on apical epithelial interactions. To circumvent these challenges, we, and others, have developed 2D human intestinal organotypic models on cell culture inserts that allow IEC polarisation and luminal accessibility [15, 16]. More recently, organ-on-chip (OoC) systems that incorporate multicellular complexity of the intestinal mucosa have also been developed and can overcome the shortcomings of 3D organoid cultures. Such intestinal OoC models have been implemented to investigate how the interplay of mouse IECs and IFs influences the cellular composition of the intestinal epithelium [17], and can model tumour growth and invasion *in vitro* [18]. However, several barriers such as high costs, lack of standardisation, and a steep learning curve limit the widespread use of these OoC systems [19, 20].

To address these limitations, we developed a human intestinal model that recapitulates the cellular complexity of the intestinal mucosa in a readily accessible and easy-to-use cell culture insert system. Specifically, we optimised and established co-culture models of adult and foetal cells to mimic IEC-IF, IEC-APC, and IEC-IF-APC interactions *in vitro*. These models were thoroughly characterised and their utility demonstrated for various applications, including studies on intestinal barrier development and function, inflammation, and viral infections.

## 2. Results

### 2.1. Epithelial-Mesenchymal Crosstalk Regulates ECM Organization and Intestinal Barrier Homeostasis

First, we developed a co-culture model that contained human IECs and IFs of adult (a-IECs and a-IFs) and foetal (f-IECs and f-IFs) origin (**Figure 1** and **Supplementary Figure 1**). Prior to co-culturing, IECs and IFs were cultured in different media formulations and characterised to confirm the retention of their specific phenotype under the new conditions optimised for the model, referred to as gut mucosa medium (GMM). The characterization of 3D organoid cultures and IEC monolayers has already been presented in our previous work [15, 16]. We performed a similar phenotypic characterization of the IF cultures and confirmed that both a- IFs and f-IFs retain phenotype and ECM deposition capacity in GMM (**Supplementary Figure 2**). Once these qualitative controls were passed, IFs and IECs were seeded sequentially and in direct contact, on a standard cell culture insert and maintained in GMM (**Figure 1A**).

**Figure 1.**
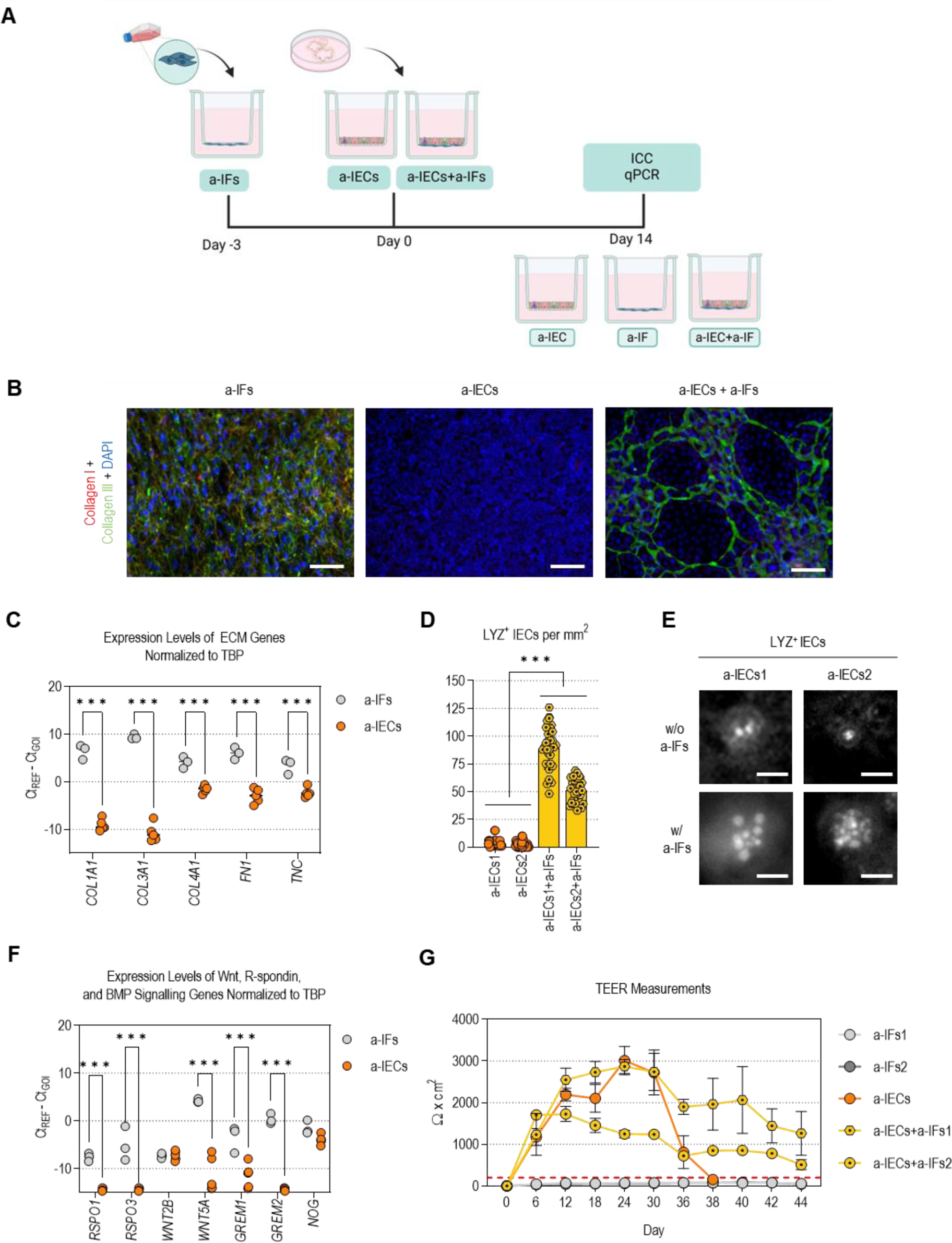
The epithelial-mesenchymal crosstalk regulates ECM organisation and intestinal barrier homeostasis. (A) Schematic representation of the workflow to establish a-IEC and a-IF co-cultures, along with their respective monoculture controls. **(B)** Representative immunofluorescence images of collagen type I and collagen type III deposition at day 14 in a-IFs (n = 2 donors), a-IECs (n = 2 donors), and a-IECs + a-IFs (2 a-IECs donors combined with 2 a-IFs donors in all possible permutations, n = 4) maintained in GMM (scale bars = 100 µm). **(C)** RT-qPCR analysis of ECM genes performed on a-IECs and a-IF monocultures maintained for 14 days on cell culture inserts in GMM. Gene expression levels are plotted as -ΔCt values. Each point represents a biological replicate (a-IFs donors: n = 3; a-IECs donors: n = 5), grey and orange colours highlight data related to a-IFs and a-IECs, respectively. Gene names are listed in the x-axis. Statistical analysis employed a two-way ANOVA, along with Šídák’s multiple comparison test. **(D)** Bar graph summarising the results of the Paneth cells quantification based on LYZ immunostaining. The bars represent the mean and standard deviation. For each sample, at least 30 images at 10X magnification per insert were analysed. Adult IECs derived from different donors are indicated with distinct numerical suffixes (e.g. a-IECs1 and a-IECs2). Statistical analysis employed a one-way ANOVA, along with Tukey’s multiple comparison test. **(E)** Representative immunofluorescence images of LYZ expression in Paneth cells at day 14 in a-IECs and a-IECs + a-IFs maintained in GMM (scale bars = 10 µm). Adult IECs derived from different donors are indicated with distinct numerical suffixes (e.g. a-IECs1 and a-IECs2). **(F)** RT-qPCR analysis of WNT, R-Spondin, and BMP signalling genes performed on a-IECs and a-IFs cultures maintained for 14 days on cell culture inserts in GMM. Gene expression levels are plotted as -ΔCt values. Each point represents a biological replicate (a-IFs donors: n = 3; a-IECs donors: n = 5), grey and orange colours highlight data related to a-IFs and a-IECs, respectively. Gene names are listed at the bottom of the plot. Statistical analysis employed a two-way ANOVA, along with Šídák’s multiple comparison test. **(G)** TEER measurements performed on a-IFs (n = 2 donors), a-IECs (n = 1), and a-IECs + a-IFs (1 a-IECs donor combined with 2 a-IFs donors, n = 2) maintained in the same medium. TEER measurements were normalised to the surface area of the cell culture inserts (0.332 cm^2^) and plotted as Ω·cm^2^ (y-axis) over time (x-axis). Each point represents the mean of at least two measurements performed on different inserts per condition, and the bars indicate the standard deviation of these measurements. Grey, orange, and yellow colours represent data pertaining to a-IFs, a-IECs and a-IECs + a- IFs. Adult IFs derived from different donors are indicated with distinct numerical suffixes (e.g. a-IFs1 and a-IFs2).

As the IFs play a key role in ECM synthesis and organisation, we assessed the effects of the epithelial-mesenchymal crosstalk on the ECM microenvironment. The deposition of collagen type I and collagen type III, the predominant collagens in the intestinal lamina propria [21], was assessed by immunostaining (**Figure 1B**). Collagen type I and collagen type III deposition was observed in the conditions that included IFs, but not in the IEC monocultures. In addition, only the co-cultures (a-IECs + a-IFs) showed the formation of a network of collagen bundles (**Figure 1B**). Gene expression analysis, performed on IEC and IF monocultures, corroborated the immunostaining data, revealing that both adult and foetal IFs contributed significantly (p < 0.001) to the ECM with higher levels of ECM-related genes (*COL1A1*, *COL3A1*, *COL4A1*, *FN1*, and *TNC*), compared to the respective IEC monocultures (**Figure 1C, Supplementary Figure 1A**). Furthermore, direct contact of IECs and IFs was necessary for the formation of the ECM network as this was not formed when the IECs and IFs were cultured on opposite sides of the culture insert (**Supplementary Figure 1B**).

In addition to their role in ECM deposition, IFs are implicated in the formation of intestinal stem cell niches and are known to secrete factors that promote epithelial proliferation and differentiation into various cell lineages [22]. Given that Paneth cells are an integral part of the stem cell niches and are important for the stability and longevity of the gut mucosa [23], we quantified the number of Paneth cells in our IEC-IF co-culture model. On day 14, the number of Paneth cells were determined by counting the number of Lysozyme (LYZ) positive cells in 2 distinct IEC-IF co-cultures (a-IECs1 + IFs and a-IECs2 + IFs) and IEC monocultures of adult origin (**Figure 1D** and **1E**). Significantly (p < 0.001) higher number of Paneth cells were observed in the IEC-IF co-cultures (89.4 ± 19.1 LYZ^+^ cells/mm^2^ in a-IECs1 + IFs and 51.1 ± 10.4 LYZ^+^ cells/mm^2^ in a-IECs2 + IFs) compared to IEC monocultures (3.8 ± 3.3 cells/mm^2^ in a-IECs1 and 2.1 ± 2.8 cells/mm^2^ in a-IECs2) (**Figure 1D**). Notably, while Paneth cells in IEC monolayers were difficult to detect due to their small size and low granularity, the LYZ^+^ granules of Paneth cells in IEC-IF co-cultures appeared larger and more numerous (**Figure 1E**). RT-qPCR analysis of the genes involved in WNT, R-Spondin, and BMP signalling, performed on monocultures, revealed that the increased presence of Paneth cell is likely supported by the IFs expressing key markers related to these signalling pathways. Significantly higher levels *RSPO1*, *RSPO3*, *WNT5A*, *GREM1*, and *GREM2* transcripts were detected in both a-IFs and f-IFs, compared to a-IECs and f-IECs respectively (**Figure 1F, Supplementary Figure 1C**). Consistent with the requirement for direct IEC-IF contact for ECM formation, the increase in Paneth cell count also correlated with direct IEC-IF co-culture. Although the administration of conditioned GMM from IF cultures (Cond. GMM) to a-IEC monolayers resulted in a significant (p = 0.04) increase in the number of LYZ^+^ cells (16.5 ± 9.5 and 10.3 ± 2.8 cells/mm^2^) compared to a-IEC monolayers, this increase was still significantly (p < 0.0001) lower than that observed in direct IEC-IF co-cultures (**Supplementary Figure 1D**).

To determine whether the increased Paneth cell count enhanced the longevity of our cultures, we evaluated long-term cell survival by measuring trans-epithelial electrical resistance (TEER) every two days for six weeks (**Figure 1G** and **Supplementary Figure 1E**). At early time points, up to day 12, the presence of IFs in the co-culture did not significantly influence the barrier function of the gut mucosa model. However, in the long term, IFs appeared to enhance barrier stability. While the barrier integrity of IEC monolayers begins to deteriorate after day 30, IEC-IF co-cultures maintain TEER values above 200 Ω·cm² for a much longer period, up to day 44. Further, as Paneth cells are primarily found in the small intestine, but are rare in the colon and rectum [24], we investigated whether the ability of IFs to induce Paneth cells was restricted by the segment from which the IECs were isolated. To test this, we established colon-derived IEC-IF co-cultures (aC-IEFs + aC-IFs) and quantified the number of LYZ^+^ cells by immunostaining (**Supplementary Figure 1F and 1G**). Unlike the increase of Paneth cell numbers in the small intestine-derived IEC-IF co-cultures, colon-derived IEC-IF co-cultures resulted in a small, but significant (p < 0.001), decrease in the number of Paneth cells (5.7 ± 2.6 and 8.0 ± 4.2 cells/mm^2^ in aC-IECs, 1.5 ± 1.3 and 1.3 ± 1.1 cells/mm^2^ in aC-IECs + aC- IFs). These findings indicate the recapitulation of segment-specific differences in the epithelial-mesenchymal interactions in a simple co-culture system.

Collectively, these results highlight the critical roles of epithelial-mesenchymal crosstalk in orchestrating ECM synthesis and organisation, as well as in modulating intestinal epithelial barrier homeostasis.

### 2.2. Epithelial-mesenchymal crosstalk in intestinal health and disease

IFs perform several immunological functions under both homeostatic and inflammatory conditions [25]. Thus, to explore their immunomodulatory properties in our model, we conducted a gene expression analysis to assess the contribution of IFs and IECs to the transcription of pro-inflammatory cytokines’ genes (**Figure 2A**). RT-qPCR analysis, performed on monolayers of IFs and IECs cultured in GMM, revealed that IECs expressed higher levels of *TNF* compared to IFs, which aligns with the function of TNFα in regulating epithelial cell turnover [26]. In contrast, no differences were noted in the expression levels of *IL-8* and *IL- 33*, while IFs expressed significantly (p < 0.001) higher levels of *IL-1β* and *IL-6*. To further investigate this at the protein level, we quantified the levels of secreted IL-6 in the supernatant of the different monocultures (**Figure 2B**). Consistent with the RT-qPCR data, high levels of IL-6 (75.2 ng/ml ± 12.2 ng/ml) were measured in supernatants isolated from a- IFs cultures. Interestingly, the levels of secreted IL-6 in a-IECs + a-IFs (2.12 ng/ml ± 1.5 ng/ml) were significantly lower (p < 0.001) compared to a-IF monocultures, suggesting that IEC-IF crosstalk may regulate IL-6 cytokine levels in the extracellular environment.

**Figure 2.**
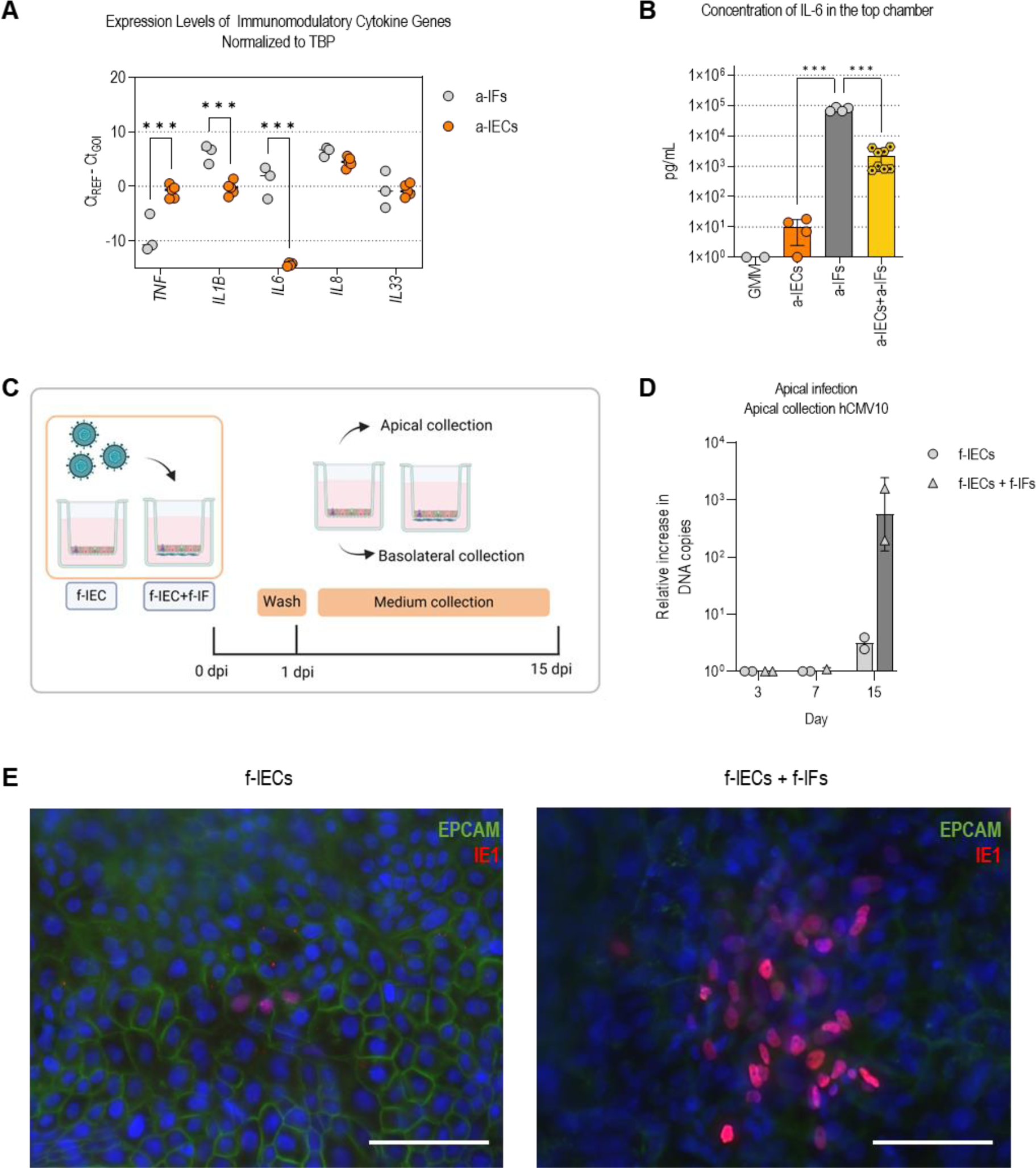
Epithelial-mesenchymal crosstalk in intestinal health and disease. (A) RT-qPCR analysis of immunomodulatory cytokine genes performed on a-IECs and a-IFs cultures maintained for 14 days on cell culture inserts in GMM. Gene expression levels are plotted as -ΔCt values. Each point represents a biological replicate (a-IFs donors: n = 3; a-IECs donors: n = 5), grey and orange colours highlight data related to a-IFs and a-IECs, respectively. Gene names are listed at the bottom of the plot. Statistical analysis employed a two-way ANOVA, along with Šídák’s multiple comparison test. **(B)** Bar graphs indicating the concentration of IL-6 secreted in the cell culture medium (GMM) by a-IF (n = 2 donors), a-IECs (n = 2 donors), and a-IECs + a-IFs (2 a-IECs donors combined with 2 a-IFs donors in all possible permutations, n = 4) after 48 hours. Each point indicates a measurement performed on different inserts, and bars represent the mean and standard deviation. Statistical analysis employed a one-way ANOVA, along with Tukey’s multiple comparison test. **(C)** Schematic representation of viral inoculation and readouts. **(D)** Average relative increase in hCMV10 DNA copy numbers over time of two independent experiments with one biological replicate (n = 1; error bars = SD). f-IEC monocultures and f-IECs + f-IFs are represented by circular and triangular grey symbols respectively. **(E)** Representative immunofluorescence images of two independent experiments on one biological replicate (n = 1) at day 15 displaying the epithelial marker EPCAM in green and hCMV viral particles by IE1 in red (scale bars = 100 µm).

Mesenchymal cells can be primary targets for some viruses, such as human cytomegalovirus (hCMV), that exhibit a broad cellular tropism [27]. As the gastrointestinal tract is a key route of infection in the case of congenital CMV infections [28], we assessed whether our new IEC- IF co-culture system could model *in vitro* hCMV intestinal infection better than IEC monocultures. To this end, we inoculated IEC monocultures and IEC-IF co-cultures of foetal origin with a clinical hCMV isolate (hCMV10). Viral replication was assessed in both f-IEC and f-IECs + f-IFs over a period of 15 days by qPCR and immunostaining (**Figure 2C**). Although infection of IECs by mouse CMV has been previously demonstrated [29], we did not observe an increase in human CMV DNA copies in the f-IEC monoculture (**Figure 2D**). In contrast, an apparent increase in hCMV10 DNA copies was measured on day 15 in f-IECs + f-IFs indicating that the IFs could be a primary target of hCMV infection in the human intestinal tract. This was consistent with infection of hCMV10 in the IF only condition (data not shown). However, in the co-cultures, hCMV appears to translocate to the basal side *via* or bypassing the f-IECs to infect f-IFs. hCMV infection of f-IFs in the IEC-IF co-cultures was also confirmed by the co- localization of immediate early antigen 1 (IE1) exclusively in EPCAM negative cells located beneath the epithelium (**Figure 2E**).

Collectively, these data indicate that the IEC-IF co-culture model recapitulates key aspects of IF immunomodulation and can serve as a suitable model to study intestinal viral infections, highlighting its potential to study intestinal diseases.

### 2.3. Gut mucosa triple-culture model with IEC, IFs, and DCs

As IL-6 signalling was observed to be finely regulated in the IEC-IF co-cultures and given the well-established role of IL-6 signalling in modulating intestinal DC function [30], we aimed to establish the first layers of the gut mucosa immune response by incorporating DCs into our IEC-IF *in vitro* model. Furthermore, the plasticity of DCs in their response to environmental cues along with their antigen presenting function made them a desirable choice for a model incorporating early immune responses. Therefore, to further enhance phenotypical and functional relevance of our *in vitro* model, we co-cultured a-IECs and a-IFs with DCs for 48 hours after IEC-IF co-culture establishment (day 14) (**Figure 3A**). The optimised media, GMM, used for IEC-IF co-culture was also verified to support DC phenotype and function prior to co- culture. (**Supplementary Figure 3**).

**Figure 3.**
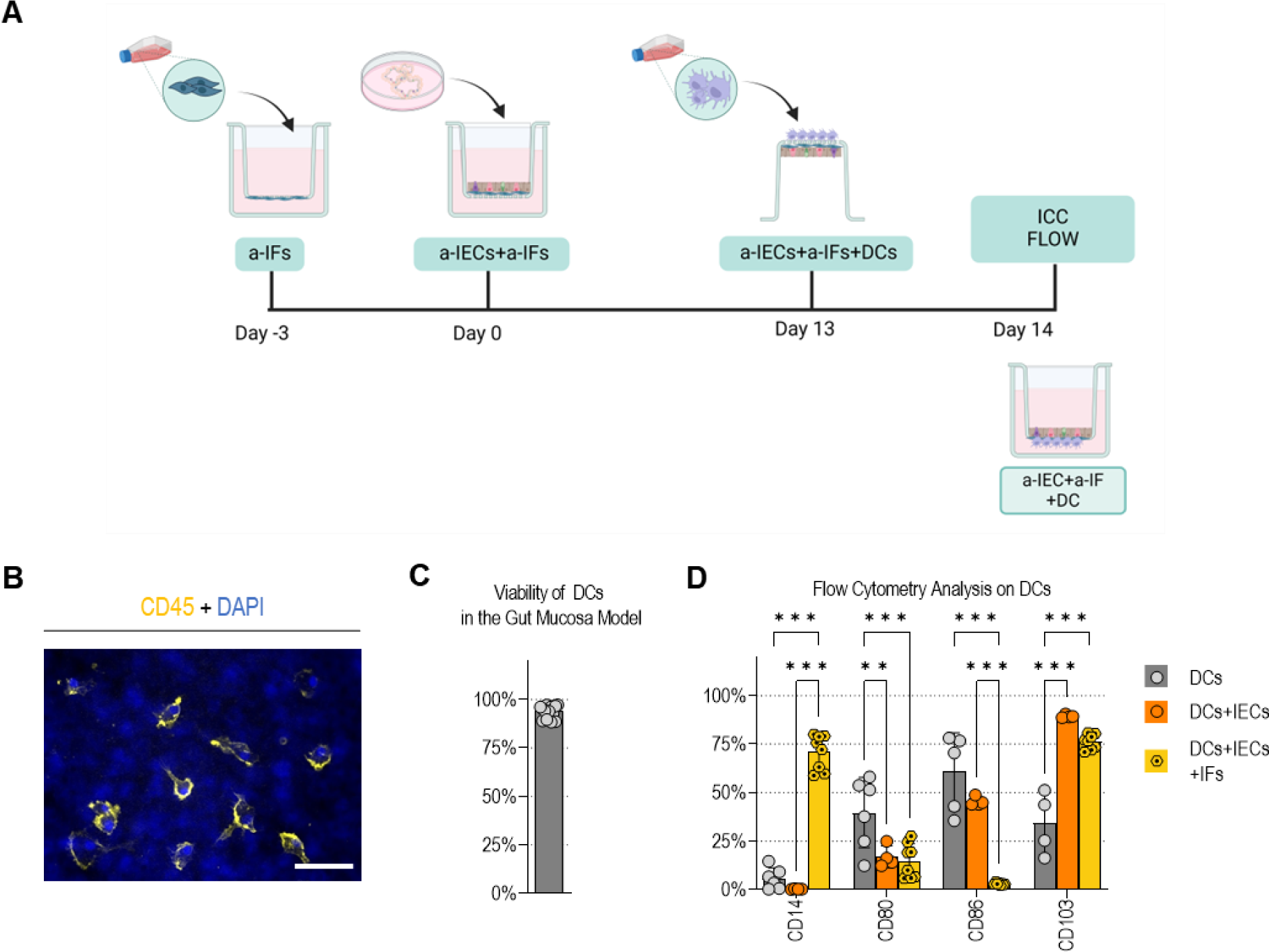
Characterization of DCs phenotype in gut mucosa triple-culture model. (A) Schematic representation of the workflow to establish a gut mucosa triple-culture model comprising a-IECs, a-IFs, and monocyte-derived DCs. **(B)** Representative immunofluorescence image at day 14 displaying CD45 expression in DCs attached to the basal side of the insert containing the IECs and IFs on the apical side (2 a-IECs donors combined with 2 a-IFs donors, and 2 monocyte-derived DCs in all possible permutations, n = 8; scale bar = 50 µm). **(C)** Cell viability assay based on acridine orange-DAPI stain performed at day 14 on DCs located in the basal chamber of the gut mucosa model maintained in GMM. Data points on the graph represent results from two independent experiments across eight biological replicates (2 a-IECs donors combined with 2 a-IFs donors, and 2 monocyte- derived DCs in all possible permutations, n = 8), with bars representing the mean and standard deviation. **(D)** Flow cytometry analysis of CD14, CD80, and CD86 expression levels at day 14 in DCs cultured in the gut mucosa model in GMM. Each point represents a biological replicate (n = 6). The bars represent the mean with the standard deviation. Statistical analysis employed a two-way ANOVA, along with Tukey’s multiple comparison test.

DC phenotype and functionality in the gut mucosa triple-culture model were thoroughly evaluated by immunostaining, flow cytometry, and ELISA (**Figure 3B, 3C, and 3D)**. Immunofluorescence microscopy verified the attachment of DCs to the basal side of the inserts with IECs and IFs on the apical side (**Figure 3B**). The presence of DCs on the basal side was evident by the detection of CD45, a pan-hematopoietic marker [31]. Furthermore, although the DCs were derived from a different donor than IECs and IFs, they were not activated as demonstrated by a branched morphology and uniform dispersion. A cell viability assay, based on acridine orange (AO)-DAPI staining, performed on DCs harvested from the basal chamber of the gut mucosa model indicated that DCs remained highly viable (93.9% ± 3.2%) (**Figure 3C**).

To elucidate the immunomodulatory effects on the DCs in the co-cultures, the expression of CD14, CD80, CD86, and CD103 was evaluated by flow cytometry for different culture conditions [DC monoculture (DCs); IEC-DC co-culture (DCs + IECs); or IEC-IF-DC triple-culture, (DCs + ECs + IFs)] (**Figure 3D**). These markers were chosen as IL-6 signalling has been previously associated with CD14 [32] expression and modulation of CD80 and CD86 expression in intestinal DCs [33] while CD103 is regarded as the main marker expressed by resident intestinal DCs [8]. The analysis revealed significant immunomodulatory effects on the DC phenotype in the triple-culture. Specifically, co-culture of DCs with IECs and IFs (DCs + IECs + IFs) resulted in a significant (p < 0.001) increase in CD14 expression and a significant (p < 0.001) decrease in CD86 expression (p < 0.004) compared to the other conditions (DCs and DCs + IECs). Additionally, CD80 expression was significantly reduced (p < 0.001) when DCs were co-cultured (DCs + IECs and DCs + IECs + IFs), compared to DC monoculture. Moreover, the expression of the intestinal DC marker, CD103, was significantly increased (p < 0.001) in the co-cultures (DCs + IECs and DCs + IECs + IFs). These data suggest that co-culturing DCs with IECs and IFs leads to phenotypic modulation, promoting the acquisition of resident intestinal tolerogenic DC markers.

Overall, these results demonstrate the possibility of incorporating several layers of cellular complexity within the gut mucosa by co-culturing intestinal epithelial and mesenchymal cells with antigen-presenting dendritic cells. Furthermore, our data indicates that the co-culture of peripheral blood-derived DCs within the IEC and IF microenvironment drives them towards an intestinal lineage enabling the study of their plasticity under different physiological conditions.

### 2.4. Modelling intestinal diseases using a complex gut mucosa triple-culture

In steady state conditions, the DCs were unaffected when cultured with allogenic IECs and IFs. To verify that DCs could maintain their function also in the gut mucosa co-cultures, we biochemically induced inflammatory conditions for 48 hours, using a commercially available cocktail of pro-inflammatory stimuli, and observed the response of the DCs (**Figure 4A**). Despite a slight, non-significant decrease in cell viability, DCs remained highly viable (89.9% ± 6.0%) even in the inflammatory cultures (**Figure 4B**). In contrast to the steady state condition, the inflammation environment triggered the formation of tightly packed clusters of DCs at the basal side of the culture insert (**Figure 4C**). This change in cellular morphology and phenotype strongly suggested that DCs underwent immune activation. Flow cytometry analysis of surface marker expression and quantification of cytokine secretion, such as IL- 12p40, serve as the primary readouts to assess DC’s activation [34]. To correlate the observed morphological changes with DC’s activation, we assessed the expression of T-cell stimulatory receptors CD80 and CD86 [35], DC’s maturation markers CD83 and CD108 [36, 37], and CD103, a marker distinguishing tolerogenic intestinal DCs promoting homeostasis, from inflammatory DCs driving effector immune responses [38]. These markers were analysed by flow cytometry (**Figure 4D**) and complemented by quantifying IL-12p40 levels in the supernatant (**Figure 4E**). Compared to steady state cultures, DCs exhibited a significant (p < 0.0001) increase in the expression levels of CD80, CD83, CD86, and CD108 in response to pro-inflammatory stimuli. Concurrently, there was a significant (p < 0.0001) decrease in CD103 expression, indicating the acquisition of an activated intestinal DC phenotype capable of initiating an adaptive immune response. Lastly, the levels of IL-12p40, detected in the supernatant via ELISA, were significantly (p value < 0.0001) higher in inflammation (1337.6 pg ± 797.5 pg) compared to steady state cultures (4.5 pg ± 0.5 pg). These results indicate that our complex gut mucosa model supports the culture of functional DCs that retain the capacity to trigger an effective immune response when exposed to pro-inflammatory stimuli *in vitro*.

**Figure 4.**
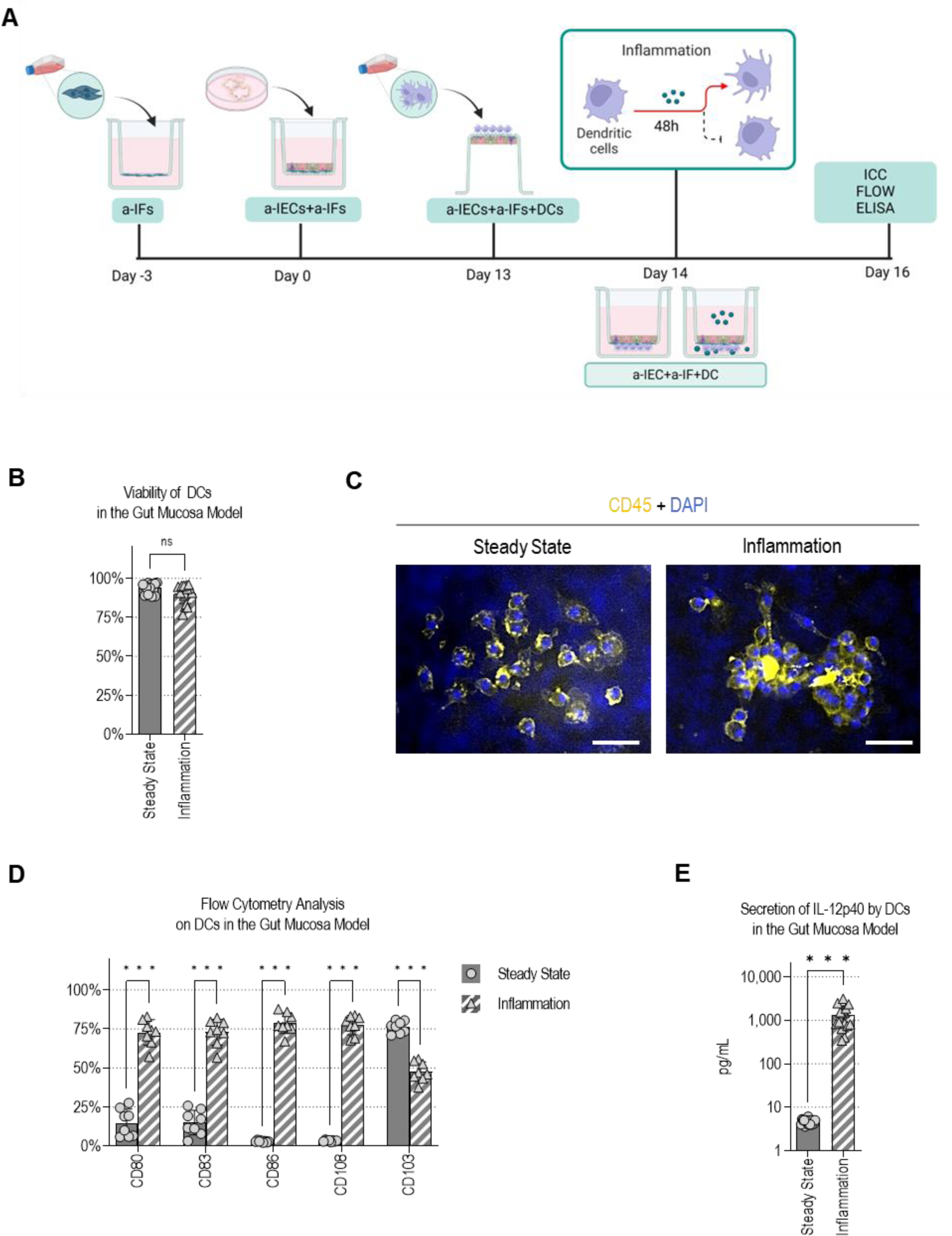
DCs remain functional in the gut mucosa triple-co-culture model. (A) Schematic representation of the workflow for modelling inflammation in the gut mucosa using the triple-culture model with dendritic cells. **(B)** Cell viability assay based on acridine orange-DAPI stain performed at day 14 on DCs located in the basal chamber of the gut mucosa model maintained in steady state or inflammation conditions. Data points on the graph represent results from two independent experiments across eight biological replicates (n = 8), with bars representing the mean and standard deviation. Statistical analysis employed a paired t-test. **(C)** Representative immunofluorescence images at day 14 displaying CD45 expression in DCs attached to the basal side of the insert containing the gut mucosa model, comparing steady state and inflammation conditions (scale bars = 50 µm). **(D)** Flow cytometry analysis of CD80, CD83, CD86, CD108, and CD103 expression levels at day 14 comparing steady state and inflammation conditions. Each point represents a biological replicate (2 a-IECs donors combined with 2 a-IFs donors, and 2 monocyte-derived DCs in all possible permutations, n = 8). Statistical analysis employed two-way ANOVA, along with Šídák’s multiple comparison test. **(E)** Bar graphs indicating the concentration of IL- 12p40 secreted in the cell culture medium by DCs in the gut mucosa model at day 14, comparing steady state and inflammation conditions. Data points on the graph represent results from two independent experiments across 8 biological replicates (n = 8), with bars representing the mean and standard deviation. Statistical analysis employed a paired t-test.

Once the functionality of the DCs in the triple-cultures were confirmed, we proceeded to assess their role in intestinal infection. To this end, we used Enterovirus A71 (EV-A71), a common intestinal pathogen. Previously, we have shown that EV-A71 can infect IECs *in vitro*, with a higher infection efficiency when the virus was inoculated from the basal lateral side of the culture insert, compared to the apical side targeted by the virus *in vivo* [39]. In parallel, we have also demonstrated that DCs are capable of transmitting EV-A71 to target cells, independent of viral replication [40]. Thus, based on these previous works, we hypothesised that EV-A71 infection of the intestinal mucosa could be better recapitulated *in vitro* by a complex culture system that contains DCs. To test this hypothesis, we took advantage of the modularity of our culture system and infected a variety of cultures [foetal IEC monoculture (f- IECs); foetal IEC - foetal IF co-culture (f-IECs + f-IFs); foetal IEC - DC co-culture (f-IECs + DCs) and foetal IEC - foetal IF - DC culture (f-IECs + f-IFs + DCs)] with EV-A71 from the apical side (**Figure 5A**). We observed a significant increase in EV-A71 viral copies by RT-qPCR and a significant increase in infectious virus particles, in all the conditions that included DCs (f-IECs + DCs and f-IECs + f-IFs + DCs), compared to conditions that did not (f-IECs and f-IECs + f-IFs) (**Figure 5B** and **Supplementary Figure 4A**). Consistent with our previous reports, viral shedding was observed only on the apical side with no viral RNA detected on the basal compartment. To rule out the increase in viral copies was due to higher cell numbers, we also infected these cultures with Echovirus 11 which infects IECs *in vitro* upon apical infection. For Echovirus 11, only a modest increase in copy numbers was detected in all the conditions including DCs (f-IECs + DCs and f-IECs + f-IFs + DCs), compared to conditions that did not (f- IECs and f-IECs + f-IFs). Interestingly only the triple-cultures (f-IECs + f-IFs + DCs) showed statistically significant (p = 0.02) differences to the f-IECs (**Supplementary Figure 4B**). Finally, as an increase in metabolic activity has been associated with DC activation following pathogen recognition [41, 42], we measured the percentage of intracellular ATP in our cultures 72 hours post infection. Again, we observed a significant increase in metabolic activity following EV- A71 infection in all the conditions that included DCs (f-IECs + DCs and f-IECs + f-IFs + DCs), compared to the conditions that did not (f-IECs and f-IECs + f-IFs) (**Figure 5C**). These results indicate that EV-A71 infection is triggered by the presence of DCs in the culture system. No significant (p > 0.05) decrease in cell viability following EV-A71 infection in any of the conditions was observed by LDH assay (**Figure 5D**).

**Figure 5.**
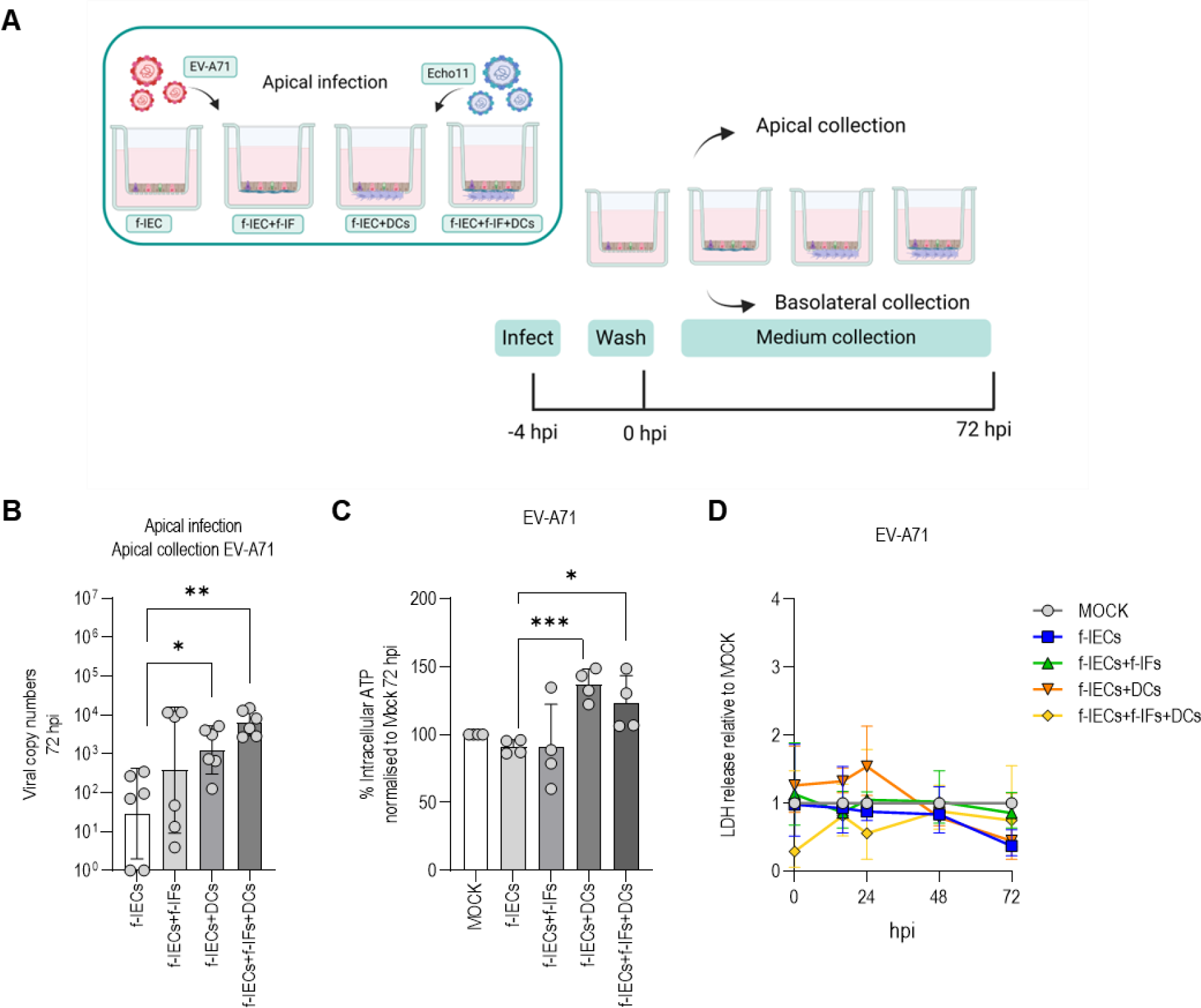
EV-A71 infection in foetal monocultures, co-cultures and gut mucosa model. (A) Schematic representation of viral inoculation and readout. **(B)** Average viral copy numbers of EV-A71, 72 hours post-infection. Data points on the graph represent results from three independent experiments across 2 biological replicates (n = 2), with bars representing the mean and standard deviation. Statistical analysis employed multiple t tests along with Bonferroni-Dunn multiple comparison tests. **(C)** Average increase (%) of intracellular ATP indicative of metabolic activity. Data points on the graph represent results from two independent experiments across 2 biological replicates (n = 2), with bars representing the mean and standard deviation. Statistical analysis employed multiple t tests along with Bonferroni-Dunn multiple comparison tests. **(D)** LDH release relative to the uninfected mock condition for all models over time. Data points on the graph represent results from two independent experiments across 2 biological replicates (n = 2), with error bars representing the standard deviation. Statistical analysis employed multiple t tests along with Bonferroni-Dunn multiple comparison tests.

In summary, the inclusion of DCs within the intestinal *in vitro* model seems to effectively mimic the dynamic interplay between DCs and their microenvironment, providing valuable insights into their functional plasticity under different physiological conditions. Additionally, the presence of DCs in our gut mucosa co-cultures significantly enhanced EV-A71 infection of IECs, highlighting the critical role of DCs in facilitating viral infection and their impact on intestinal immune responses.

### 2.5. Gut mucosa triple-culture model with IEC, IFs, and macrophages

In the previous sections, we focused on the addition of DCs to generate an intestinal mucosa model. However, DCs are not the sole APCs in the intestinal mucosa, and macrophages (MΦ) play pivotal roles in maintaining intestinal homeostasis, as well as in modulation of inflammation, wound healing, infection, and tumorigenesis [43]. Therefore, to create an advanced gut mucosa model that could recapitulate macrophage plasticity and functions *in vitro*, we also optimised conditions to establish triple-cultures comprising a-IECs, a-IFs, and monocyte-derived MΦ. Distinct macrophage subtypes (M1 and M2) were generated during addition of non-activated MΦ (M0) to a-IECs + a-IFs co-cultures through the administration of specific activation factors to steady state culture conditions (GMM) for 48 hours (**Figure 6A**). The phenotype and function of the MΦ within the gut mucosa model were thoroughly assessed using cell viability, flow cytometry, and immunostaining. First, we used immunofluorescence microscopy to verify the presence of MΦ attached to the basal side of the culture insert (**Figure 6B**). Immunostaining for the pan-hematopoietic marker CD45 confirmed the presence of MΦ attached to gut mucosa inserts in all culture conditions. The cell viability assay based on AO-DAPI staining performed on MΦ harvested from these inserts indicated that every macrophage subtype remained highly viable (93.7% ± 4.0% M0; 95.4% ± 3.2% M1; and 94.7% ± 4.4% M2) during co-culture with IECs and IFs (**Figure 6C**).

**Figure 6.**
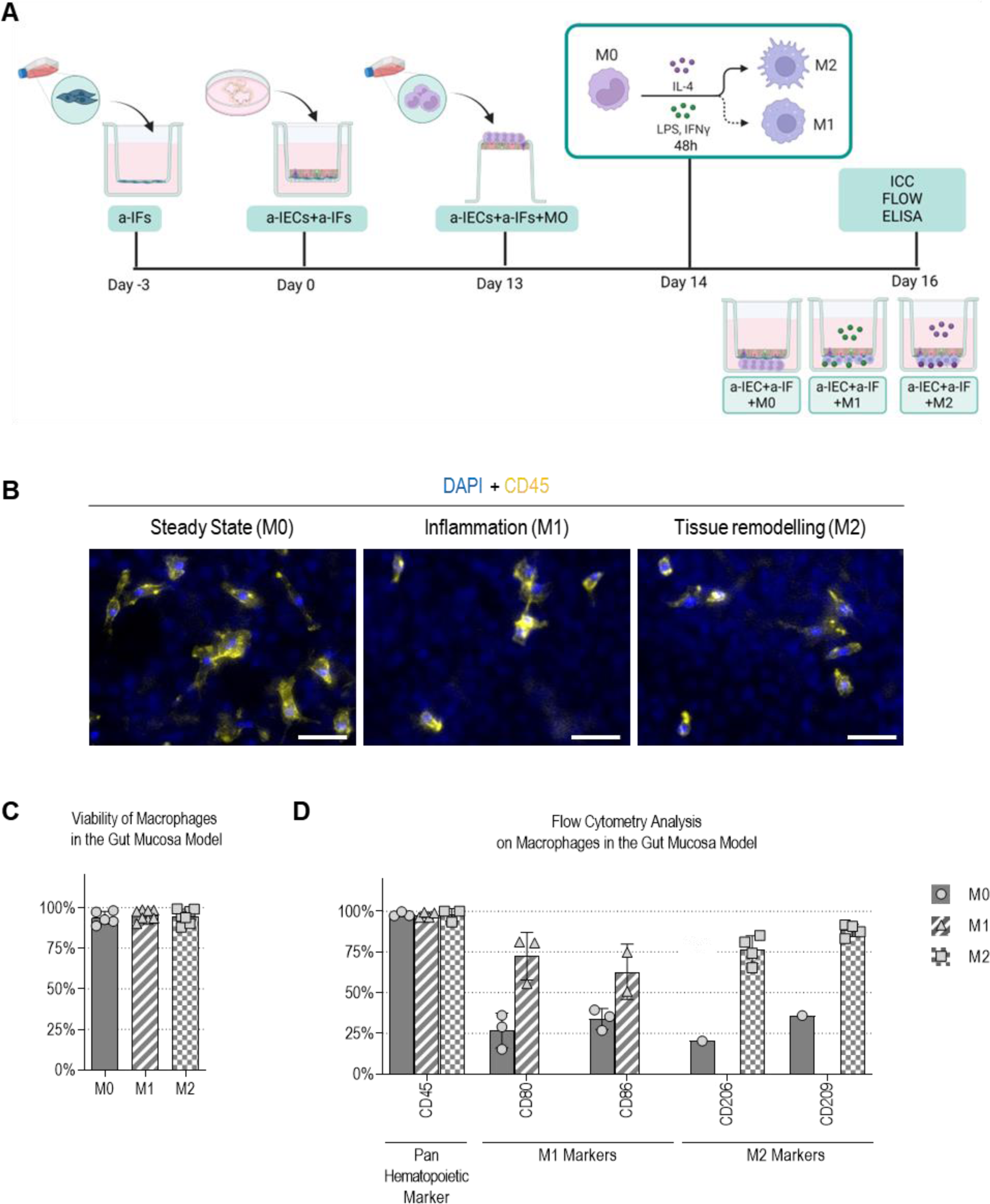
Macrophages in the gut mucosa model maintain viability, phenotype, and functionality. (A) Schematic representation of the workflow for modelling inflammation in the gut mucosa using the triple-culture model with MΦ. **(B)** Representative immunofluorescence images at day 14 displaying DAPI and CD45 expression in MΦ attached to the basal side of the inserts containing the gut mucosa triple-culture, maintained in GMM, either with or without supplementation of M1 or M2 activation factors (n = 8; scale bars = 50 µm). **(C)** Cell viability assay based on acridine orange-DAPI stain performed at day 14 on MΦ located in the basal chamber of the gut mucosa model maintained in GMM, either with or without supplementation of M1 or M2 activation factors (10 ng/mL LPS + 50 ng/mL IFN-γ, or 10 ng/mL IL-4 respectively). Data points on the graph represent data related to 2 biological replicates (n = 2) and 4 technical replicates, with bars representing the mean and standard deviation. **(D)** Flow cytometry analysis of CD45, CD80, CD86, CD206, and CD209 expression levels at day 14 in M0, M1, or M2 macrophages cultured in the gut mucosa model in GMM. Each point represents data related to MΦ in a specific gut mucosa model (n = 4). The bars represent the mean with the standard deviation. Full grey bars with square symbols indicate data related to M0 macrophages, whilst striped bars with triangle symbols indicate data relative to M1 macrophages, and chequered bars with circle symbols indicate data relative to M2 macrophages.

Phenotypical and functional characterization of MΦ activation state, induced by infections or exposure to biochemical stimuli, is commonly performed by flow cytometry analysis of surface marker expression [44–47]. M1 macrophages are identified by high levels of expression of CD80 and CD86, crucial molecules involved in antigen presentation, T cell activation, and initiation of inflammation [48]. On the other hand, M2 macrophages are characterised by high levels of expression of CD206, and CD209, and play a role in tissue repair, immunosuppression, and the resolution of inflammation [48]. In our study, flow cytometry analysis indicated that all types of MΦ constitutively expressed the pan-hematopoietic marker CD45 (**Figure 6D**). M0 macrophages in the gut mucosa model showed low levels of both M1 (26.7% ± 10.6% CD80 and 33.8% ± 6.4% CD86) and M2 (20.27% CD206 and 35.91% CD209) markers. However, when M1 activation stimuli was administered, the MΦ showed increased expression of CD80 (72.4% ± 14.6%) and CD86 (62.8% ± 17.0%). Similarly, when M2 activation stimuli were administered, the MΦ showed increased expression of CD206 (76.4% ± 8.7%) and CD209 (88.2% ± 3.8%). These data highlighted the successful activation of macrophages to the M1-like and M2-like state from M0 during co-culture in our complex gut mucosa model.

These results demonstrate that our advanced gut mucosa co-culture system effectively recapitulates the steady state, pro-inflammatory, and tissue remodelling characteristics of M0, M1, and M2 macrophages, respectively. This suggests that our model has the potential to serve as a platform for investigating macrophage behaviour, function, and interaction with other cell types in a highly physiologically relevant environment.

## 3. Discussion

Recently, OoC systems have been developed that elegantly recapitulate important features of the intestinal mucosa, but barriers such as high costs, lack of standardisation, needs of special equipment and a steep learning curve limit their widespread use. To overcome this, while balancing the simplicity of organoids and complexity of OoC systems, we assembled a complex intestinal mucosa model by reconstructing the structure and architecture of the intestine *in vivo* on widely used cell culture inserts. This resulted in establishing a new modular *in vitro* co-culture model that recapitulates the complexity of the gut mucosa including epithelial, mesenchymal, and immune cells. We optimised culture conditions and protocols to make the model robust, reproducible, and easy to use. We showed that it can be easily adjusted to comprise a variety of cell types in different combinations to adapt to specific scientific questions. We also provided evidence of the value of a primary derived human gut mucosa model to study multicellular crosstalk necessary for maintaining homeostasis of the intestinal barrier, and its role in disease, inflammation, and viral infections.

Fibroblasts in the gut are responsible for ECM deposition to support the pericellular matrix and basal membrane of epithelial cells [49]. Here, we described the importance of direct contact of IFs and IECs in an *in vitro* culture that is required to achieve levels of ECM deposition and organisation similar to *in vivo* [50, 51]. This close crosstalk also promoted the differentiation of Paneth cells which are important cellular components of the stem cell niche and contribute towards tissue regeneration [52–54]. Interestingly, in our model, Paneth cell differentiation was promoted only in the context of the small intestine, with no similar effects observed when modelling the colonic epithelium. These *in vitro* findings align with the study of Hickey and colleagues [1] who, using single cell-RNA sequencing of human tissue, showed that Paneth cells specifically populate the small intestine and organise into specialised “neighbourhoods”. They postulated that such functional neighbourhoods across the intestinal tissue define the cellular composition and function of the human gut [1]. Similarly, in an *in vivo* study performed using murine models, Maimets and colleagues [50] found that the mesenchyme plays a critical role in defining the regional identity of the intestinal epithelium by modulating the WNT signalling. The fact that such segment-specific differences can be now recapitulated in a cell culture system, such as the one described here, represents an invaluable contribution to further increase the predictive value of results obtained *in vitro*.

With the observed increase in Paneth cell numbers in our IEC-IF co-cultures, we wanted to determine whether the fibroblasts similarly engage in dynamic, bidirectional communication with neighbouring cells under homeostatic and inflammatory conditions [25]. *In vivo,* intestinal fibroblasts interact with resident immune cells through the secretion of pro-inflammatory cytokines such as TNFα, IL-1β, IL-6, IL-8, and IL-33 [56, 57]. Through the tight regulation of the levels of secretion of these cytokines, the IFs immune modulate APC function [25]. In our co-culture system, we observed how the interaction between intestinal fibroblasts and epithelial cells influences a number of these cytokines, including IL-6. This cytokine was of particular interest as it regulates epithelial proliferation, ECM remodelling, and immune cell activation [58–60]. Interestingly, IL-6 dysregulation has been linked to inflammatory bowel diseases (IBD) which is marked by chronic inflammation that fuels the development of intestinal fibrosis [61], and for this reason the cytokine has recently become a therapeutic target for intestinal pathologies [62–65]. The addition of mesenchymal cells combined with the epithelial barrier enables the potential study of fibrosis, while current modes solely focus on epithelial barrier disruption induced by biochemical pro-inflammatory stimuli [66].

Through the establishment of this complex *in vitro* model, we could successfully bio mimic the influence of the crosstalk between the different cell types and the environment has on cell identity. In particular, dendritic cells demonstrated high sensitivity to these factors that have significant effects on cytokine secretion. In addition, DCs acquired a phenotype that closely resembles resident intestinal tolerogenic DCs both in homeostatic [67, 68] and inflammatory conditions [69, 70] described *in vivo*. With the addition of macrophages to our co-culture model, we were also able to effectively replicate the steady state, pro-inflammatory, and tissue remodelling characteristics of M0, M1, and M2 macrophages. These results align with previous findings using the Caco-2 cell line and Thp-1 cells [77]. However, similar to studies utilizing more physiologically relevant organoid systems [78], our optimized model more closely mimics the cellular environment found in the intestinal niche *in vivo*. This, combined with the modularity of our approach, introduces a new platform for investigating macrophage behaviour, function, and interactions with multiple cell types and environments.

As a proof of concept for the infection modelling, we tested both the IEC-IF and the triple culture mucosa model on viral infections. The added complexity of the IEC-IF culture as a result of the the close interaction established between IFs and IECs allowed us to observe an efficient replication of a clinical strain of hCMV *in vitro*. Unlike others that used unpolarised *in vitro* intestinal models with limited success in supporting effective viral replication [55], our highly polarised IF-IEC co-culture system showed high susceptibility to hCMV infection. hCMV inoculated into our model targeted the intestinal fibroblasts, similarly to *in vivo* [27], indicating the potential of this co-culture system in virology applications.

Furthermore, it is known that DCs play an essential role in triggering innate immune response to viral infections in the gut [71, 72]. Previously, we have shown the susceptibility of *in vitro* polarised intestinal monolayers to EV-A71 infection, in which viral replication was only detected upon inoculation of the virus from the basal side [39]. However, it is understood that EV-A71, similar to other picornaviruses, is transmitted through the oral-faecal route [73] and therefore, interacts with the gastrointestinal epithelium from the apical side. Our previous results alongside the known ability of DCs to transmit the virus to cells downstream [40], led us to hypothesise that a possible mechanism of EV-A71 infection could occur *via* a “trojan horse” mechanism, through the subversion of the innate immune and APC function of DCs. Infection of our gut mucosa model with EV-A71 allowed us to better mimic the viral replication happening *in vivo*, with the virus being inoculated from the apical side and the viral replication levels equalling basolateral infection [39] in the presence of DCs within the model. Interestingly, a number of other viruses, including poliovirus and norovirus, use cells to bypass the intestinal barrier and initiate infection basolaterally [74–76]. Considering this, we believe that our model holds significant potential for recapitulating the molecular mechanisms of various viruses, many of which remain poorly understood. Additionally, it can serve as a valuable platform for antiviral drug screenings.

A key question that remains to be answered is the role of the mesenchymal and immune cells in intestinal regeneration upon damage induced by inflammation and diseases. The role of epithelial-mesenchymal-immune crosstalk is well known and established in epithelial maintenance [79]. However, how epithelial identity is maintained during repair and if mesenchymal immune cell crosstalk plays a role in terms of classical homeostatic niche factors is yet to be determined [80]. We believe that having readily available *in vitro* models capable of biomimicking the epithelial-mesenchymal-immune cells crosstalk would be beneficial in the elucidation thereof, underscoring the need and the timeliness of our model.

Overall, with our modular co-culture model, we have provided an advancement on the cellular complexity over the current *in vitro* intestinal models while ensuring their accessibility and physiological relevance in a widely used culture platform, such as the cell culture inserts. We also recapitulated *in vitro,* the crosstalk between multiple cell types and the first key elements of the intestinal mucosa, including intrinsic and innate immunity. Our advanced gut mucosa model contributes towards increasing the physiological relevance of current culture models such as intestinal organoids. Furthermore, our model in combination with existing OoC models hold immense potential to advance the current state-of-the-art for studying human intestinal health and disease.

## 4. Study limitations

For this study, our focus was on introducing cellular diversity and specialisation by incorporating the first layer of the human intestinal mucosa. However, our model does not capture the entirety of the mucosal complexity and lacks other key cell populations. These include, but are not limited to, tuft cells, B-cells, and T-cells. Incorporating these cells will require even more refined culturing conditions and a completely isogenic system. Another key component of the intestinal mucosa that is currently lacking in our model is the microbiome. However, we, and others, have demonstrated the possibility of incorporating commensals in organotypic models and similar approaches will further expand the physiological relevance of intestinal mucosa models [81–83].

## 5. Materials & Methods

### 5.1. Cells and equipment

Unless otherwise stated, all cells, media, media supplements and reagents were provided by STEMCELL Technologies (Canada); plasticware was purchased from Starlab (UK); 6.5 mm cell culture inserts made of polyethylene terephthalate (PET), with a pore size of either 0.4 µm or 3 µm, and purchased from CellQART (Germany) or Greiner Bio-One (UK), and qPCR primers were purchased from Integrated DNA Technologies (IDT, Belgium). Colon fibroblasts (ATCC, cat# CRL-1459) and small intestinal fibroblasts (Innoprot, cat# P10760) were purchased from ATCC (USA) and 2B Scientific (UK) respectively.

### 5.2. Cell isolation

#### 5.2.1. Ethics statement

Human foetal intestinal tissue, gestational age 16–18 weeks, was obtained from a private clinic by the HIS Mouse Facility of the Amsterdam University Medical Center. All donors supplied written informed consents for the use of foetal material for research purposes. These consent forms are kept at the clinic and the information available to the Amsterdam UMC does not allow identification of the donor without disproportionate efforts. The use of the anonymised material for medical research purposes is covered by Dutch law (Wet foetaal weefsel and Article 467 of Behandelingsovereenkomst).

#### 5.2.2. Generation of human foetal intestinal organoid cultures

For the generation of foetal small intestinal organoids, crypts were isolated from foetal intestinal tissue as described previously [15]. Isolated crypts were suspended in Matrigel, dispensed in three 10 μL droplets per well in a 24-well tissue culture plate and covered with 500 μL medium. Enteroid cultures were routinely maintained at 37°C, 5% CO2, and 95% humidity in Human IntestiCult™ Organoid Growth Medium Human (IntestiCult OGMh) supplemented with 100 U/mL penicillin/streptomycin (Pen-Strep) (Lonza). Medium was replenished every second day, and organoids passaged every 3-5 days.

#### 5.2.3. Generation and maintenance of human foetal intestinal fibroblast cultures

Following crypt isolation, tissue pieces were further digested in 1mg/mL Collagenase I (cat #100-0677) for 45 min at 37ᵒC. Collagenase activity was inhibited by 0.05 mM EDTA, and the resulting cell suspension was passed through a 70 µm cell strainer. The collected cell suspension was centrifuged at 300 x g for 5 min, the supernatant discarded, and the cells resuspended in advanced DMEM/F12 (AdvDMEM/F12) (Gibco) supplemented with 100 U/mL (v/v) Pen-Strep, 1% (v/v) Glutamax and 2% (v/v) heat-inactivated foetal bovine serum (FBS, Sigma-Aldrich) (2% AdvDMEM/F12). The Primary human intestinal fibroblasts were allowed to adhere to the culture flask overnight and the medium was replenished thereafter. Intestinal fibroblasts were then routinely cultured at 37ᵒC, 5% CO2 and 95% humidity in 2% AdvDMEM/F12. The culture medium was replenished every second day and cells were subcultured by trypsinization upon reaching 70% confluence.

5.3. *In vitro* cultures

#### 5.3.1. Maintenance of adult human intestinal fibroblasts

Intestinal fibroblasts (IFs) derived from small intestine were purchased from 2B Scientific (cat #P10760). Colon fibroblasts were purchased from ATCC (cat #CRL-1459). IFs were expanded using the MesenCult™ - ACF Plus Culture Kit (cat #05448) that includes Animal Component-Free Cell Attachment Substrate (ACF-CAS) used for coating standard cell culture-treated flasks. Cryopreserved IFs were plated at a density of 1 x 10^3 cells/cm² in complete MesenCult™ - ACF Plus Medium (cat #05445) and incubated at 37°C until 90% confluent. The culture medium was replenished every second day and cells were subcultured by trypsinization upon reaching 70% confluence.

#### 5.3.2. Maintenance of adult and foetal human intestinal organoids

Intestinal organoid cultures were maintained and expanded using IntestiCult OGMh in 10 µL Matrigel® as previously described [39], and according to the manufacturing protocol. The medium was changed every other day, and the organoids were passaged every 7 days.

#### 5.3.3. Differentiation of monocytes into dendritic cells

Human Peripheral Blood Monocytes, either cryopreserved (cat #70034) or freshly isolated Human Peripheral Blood Leukopak (#70500), were differentiated into dendritic cells (DCs) using ImmunoCult™ Dendritic Cell Culture Kit (ImmunoCult DC), following the manufacturer’s instructions. Monocytes were isolated from Human Peripheral Blood Leukopak using EasySep™ Human CD14 Positive Selection Kit II (cat #17858).

#### 5.3.4. Differentiation of monocytes into macrophages

Human monocytes, either cryopreserved or freshly isolated from whole blood, were differentiated into macrophages using ImmunoCult™ SF Macrophage Differentiation Medium (ImmunoCult MΦ), following the manufacturer’s instructions.

#### 5.3.5. Establishment of the advanced gut mucosa model comprising intestinal epithelial cells, intestinal fibroblasts, and antigen-presenting cells

The gut mucosa model, which includes IECs, IFs, and APCs, is established using a modular workflow, where each cell type is sequentially added to standard cell culture inserts. Alternatively, intermediate models comprising only two cell types (IEC-IF or IEC-APC) were also developed. All models were established using a single co-culture medium, IntestiCult™ Organoid Differentiation Medium Human (IntestiCult ODMh), referred to as GMM, from the moment the cells are seeded on the inserts. Briefly, on day -3, cell culture inserts were coated with 100 µL ACF-CAS for 1-hour, followed by seeding either adult or foetal IFs at a density of 1 x 10^5/cm^2^ in 300 µL IntestiCult ODMh in the apical chamber of the cell culture inserts. To the basal chamber 700 µL IntestiCult ODMh and IFs incubated at 37°C until confluent. On day 0, adult or foetal intestinal organoids were mechanically and enzymatically fragmented into a single-cell suspension. The resulting IECs were then plated at a density of 1 x 10^6/cm² onto the confluent adult or foetal IFs in the apical chamber of cell culture inserts. IEC-IF co-cultures were maintained in IntestiCult ODMh. Medium was changed daily in the apical chamber, and every other day in the basal chamber. On day 14, APCs were harvested, inserts containing IEC-IF co-cultures were prepared by removing the medium and inverting them on an extra- depth plate. Subsequently, APCs were seeded onto the basal side of these inserts at a density of 1 x 10^5/cm^2^ and allowed to attach at 37°C for at least 3 hours. Inserts were then returned to the upright position in tissue culture-treated plates, with 300 µL and 700 µL of IntestiCult ODMh in the apical and basal chambers, respectively. To ensure the IEC-IF co-cultures were ready for downstream applications, the integrity of the intestinal barrier was verified by (i) TEER measurements, (ii) paracellular permeability assay, and (iii) visualisation of a uniform network of tight junctions by ZO-1 immunostaining.

### 5.4. Viruses

#### 5.4.1. Propagation of Enterovirus A-71 and Echovirus 11

Human rhabdomyosarcoma (RD) cells (CCL-136^™^ ATCC) and African green monkey kidney (Vero) cells (CCL-81^™^ ATCC), were cultured in Eagle’s minimum essential medium (EMEM) (Lonza) supplemented with 8% FBS, 100 U/mL Pen-Strep, 0.1% (v/v) L-glutamine (Lonza), and 1% (v/v) non-essential amino acids (100X) (ScienceCell Research Laboratories). Cell lines were routinely cultured at 37°C with 5% CO2, 95% humidity and passaged every 7 days using 0.05% (v/v) Trypsin/EDTA (Gibco). EV-A71 C1-91-480 obtained from the RIVM (GenBank: AB552982.1) and Echo 11 was propagated in RD and Vero cells respectively.

#### 5.4.2. Propagation of human Cytomegalovirus

Human embryonic lung fibroblast (HEL) cells (isolated at Amsterdam UMC) were cultured in EMEM supplemented with 8% FBS, 100 U/mL pen-step, 0.1% (v/v) L-glutamine and 1% (v/v) non-essential amino acids at 37 °C with 5% CO2. A clinical isolate of hCMV was derived from patient urine material (isolated at Amsterdam UMC) and propagated in HEL cells and retinal pigment epithelial cells (ARPE-19). A hCMV TB40e-mCherry strain was gifted by and propagated in HEL cells.

5.5. *In vitro* modelling of viral infections

#### 5.5.1. Viral infection with EV-A71 and Echo 11

Viral infections on f-IECs, f-IECs + f-IFs, f-IECs + DCs and f-IECs + f-IFs + DCs cultures were performed as described previously [84, 85]. Briefly, viral stocks (EV-A71 and Echo 11) were diluted to 10^5^ TCID50/50 μl in IntestiCult ODMh. Cultures were apically inoculated with virus, by removing 50 μL of medium and adding 50 μL of viral inoculum. The cultures with virus were incubated for 4 h at 37°C with 5% CO2. Following incubation, unbound virus was washed away from the apical compartment with advDMEM/F12 three times. Inserts were incubated for 10 min with IntestiCult ODMh after which the 0 hours post infection (hpi) time point was collected by removing 200 μL from the apical and the basal compartment. After collection, medium was replaced apically and basally. Cultures were incubated for 72h, with sample collection both apically and basally at 16, 24, 48 and 72 hpi. Collected medium samples were stored at -70 °C until further use.

#### 5.5.2. Viral infection with hCMV

Virus stocks (clinical strain hCMV10 and TB40e-mCherry) were thawed in a water bath at 37°C and centrifuged at 3.300 x g for 10 min at RT. Apical medium was removed from f-IECs and f- IECs + f-IFs cultures and 100 µL virus stock supernatant was added to the apical side for 24 h at 37 °C, 5% CO2. After 24 h incubation, the virus containing medium was removed and inserts were washed three times with pre-warmed Advanced DMEM/F12. After washing, apical and basal medium was removed and collected for the 1-day post infection (dpi) time point. IntestiCult ODMh was added, and medium was changed and collected every other day until 15 dpi. Collected medium samples were stored at -70 °C until further use.

#### 5.5.3. Median tissue culture infectious dose

Supernatant samples from EV-A71 and Echo 11 at 0 and 72 hpi were used to determine the presence of infectious viral particles; titrations were performed on the RD cell line. Seven ten-fold dilutions of each sample were performed and 50 µL of each dilution was added to a 96- well plate, after which 200 µL of specific cells were added. Plates were incubated for seven days until scoring of the cytopathic effect (CPE) and calculation of the median tissue culture infectious dose (TCID50) according to the Reed and Muench Method [86]. Values at 72 hpi were normalized to 0 hpi to determine the increase of infectious particles.

#### 5.5.4. Viral RNA isolation and qPCR EV-A71 and Echo11

RNA was isolated from cultures infected with EV-A71 and Echo11. Media samples (25 μL) were added to 300 μL lysis buffer and the total RNA was isolated using the PureLink^TM^ RNA Mini kit (Thermo Fisher) according to the manufacturer’s instructions. Equal volumes of eluted RNA were used for reverse-transcription using the SuperScript™ II Reverse Transcriptase synthesis kit (Thermo Fisher). A volume of 5 μL of cDNA was used for quantitative PCR (qPCR) on a CFX Connect Real-Time PCR Detection System (Bio-Rad). SYBR Green qPCRs were performed using the pan entro primers 5’– GGCCCTGAATGCGGCTAAT –3’; reverse primer 5’– GGGATTGTCACCATAAGCC –3’; as described previously (Benschop et al., 2010).

#### 5.5.5. Viral DNA isolation and qPCR hCMV

Viral DNA was isolated from cultures infected with hCMV. 100 µL media samples were collected from the apical and basal chamber and viral DNA was isolated using the ISOLATE II Genomic Kit (Meridian Bioscience,). The following primers and probe were used for quantitative PCR (qPCR) on a CFX Connect Real-Time PCR Detection System: forward primer 5’– CACGGTCCCGGTTTAGCA –3’; reverse primer 5’– CGTAACGTGGACCTGACGTTT –3’; and Taqman probe FAM – TGTAACCGCGATCCTCGGGCAGATA – TAMRA.

### 5.6. *In vitro* modelling of inflammation

To simulate an *in vivo* inflammatory environment and activate specific subsets of APCs, various combinations of cytokines were supplemented into the complete or intermediate gut mucosa models. ImmunoCult™ Dendritic Cell Maturation Supplement was added to IntestiCult ODMh in both the apical and basal chambers to induce DC activation. Alternatively, 10 ng/mL LPS and 50 ng/mL IFN-γ were supplemented to IntestiCult ODMh for classical macrophage activation (M1), or 10 ng/mL IL-4 for alternative macrophage activation (M2), also in the apical and basal chambers of inserts containing the complete gut mucosa model. For long-term experiments (over 48 hours), medium changes were performed every other day with 300 µL and 700 µL in the apical and basal chambers, respectively. Each time, non-adherent APCs collected during these steps were rescued, and returned to the basal chamber.

### 5.7. Cell viability

#### 5.7.1. Quantification of intracellular adenosine triphosphate

Cell viability and cellular metabolic activity was determined based on intracellular adenosine triphosphate (ATP) production, using the 3D CellTiter-Glo Luminescent cell viability assay (Promega) as published previously (Wrzesinski et al., 2021) with minor modifications. Briefly, medium was aspirated and 100 µL PBS was added to each sample well. Cells were lysed with 100 µL 3D CellTiter-Glo lysis buffer and shaken in the dark for 30 min. Following incubation, the lysate was transferred to black clear bottom 96-well plates and the luminescence was measured in an H1 Synergy plate reader (BioTek). The data was normalized with reference to a standard curve for ATP (Sigma-Aldrich) and relevant controls.

#### 5.7.2. Extracellular lactate dehydrogenase

Cell toxicity was determined by extracellular lactate dehydrogenase (LDH) release, using the LDH-Glo™ Cytotoxicity assay (Promega) following manufacturer’s instructions. Briefly, culture medium samples (5 µL) were added to 95 µL LDH storage buffer and stored at -70°C until further processing. The assay was performed by transferring 50 µL of the sample and LDH standard in LDH storage buffer to a black clear bottom 96-well plate. LDH detection reagent (50 µL) was added to standard and sample wells and incubated at room temperature in the dark for one hour. Luminescence was measured in an H1 Synergy plate reader. Data was normalized with reference to a standard curve of LDH and relevant controls.

#### 5.7.3. Quantification of live and dead cells by automated fluorescent cell count

Cell viability was assessed using Nucleocounter NC-250 (Chemometec), an automated fluorescent cell counter. Cell cultures were first dissociated into single-cell suspensions using either ACCUTASE or TrypLE. Collected cells were centrifuged at 300 x g for 5 min and resuspended in their respective culture medium. A 20 µL aliquot of the cell suspension was stained with 1 µL of AO-DAPI Solution 18 (Chemometec) to differentiate between live and dead cells. Samples were analysed using the Nucleocounter NC-250 to estimate cell viability.

### 5.8. Trans electrical epithelial resistance measurements (TEER)

TEER was measured using the EVOM2 or EVOM3 Epithelial Voltohmmeter (World Precision Instruments), routinely calibrated with a set of standard resistors. Resistance values of a coated empty insert were used as background. All TEER measurements were performed in triplicate per donor and the average was used for calculations.

### 5.9. FITC-dextran permeability assay

Paracellular permeability was assessed using the fluorescent tracer FITC-Dextran 4 kDa (FD4). A solution of 1 mg/mL FD4 was added to the apical chamber of cell culture inserts. After incubation on a shaker at 70 rpm and 37°C for either 3 or 24 hours, 100 µL aliquots of medium were sampled twice from the basal chamber and transferred to a black flat-bottom 96-well plate. Fluorescence intensity at 520 nm was measured using either a SpectraMax® M3 multi- mode microplate reader or a Synergy hybrid microplate reader, with the excitation wavelength set at 490 nm, a cut-off at 515 nm, and an emission wavelength at 520 nm. Data was normalised to a standard curve with known concentrations of FITC-Dextran in IntestiCult ODMh. The plate blank was established using a 100 µL aliquot of IntestiCult ODMh medium. Control measurements included a loading control (1 mg/mL FD4 in IntestiCult ODMh) and an empty insert control (cell culture insert without cells) for each experiment.

### 5.10. Gene expression analysis

Total RNA was purified using the RNeasy Mini kit spin columns or the Ambion PureLink® RNA Mini kit, according to the supplier’s protocol. RNA quantification was performed using Nanodrop 2000 (ThermoFisher Scientific, USA). Reverse transcription was carried out with High Capacity cDNA Reverse Transcription Kit with Complementary DNA (cDNA) samples diluted to a final concentration of 5 ng/ml. Quantitative real-time polymerase chain reaction (RT-qPCR) assay was performed with TaqMan™ Fast Universal PCR Master Mix (2X) no AmpErase™ UNG and 0.5 µL of primer/probe set. Predesigned qPCR assays were purchased from Integrated DNA Technologies (USA) and Applied Biosystems (USA) or were custom designed using IDT PrimerQuest Tool (IDT, USA) and sequence homogeneity was confirmed by comparison to all available sequences on the GenBank database using BLAST (http://www.ncbi.nlm.nih.gov/BLAST/). The primers used in the study were purchased from IDT (Belgium) and are listed in **Supplementary Table 1**. RT-qPCR was performed using the StepOnePlus Real-Time PCR system (Life Technologies, USA). Standard curves and efficiency tests were generated for each target. Unless differently stated in the figure legends, gene expression levels are plotted as -ΔCt values, values are normalised to the average threshold cycle (Ct) values for the gene of interest (GOI) against the average Ct values for the reference genes, from 3 technical replicates.

### 5.11. Immunocytochemistry

Human foetal intestinal mucosa cultures were fixed in 4 % (v/v) paraformaldehyde (PFA) or cold methanol for 30 and 10 min respectively at RT and stored in PBS at 4 °C. To reduce autofluorescence inserts were treated with 0.3% (w/v) Sudan Black B (Sigma- Aldrich, 199664- 25G) in 70% (v/v) Ethanol for 30 min at RT. Inserts were immersed and washed five times in PBS. Overnight blocking was performed, using Fish Serum Blocking Buffer (ThermoFisher, 37527) at 4 °C. A Liquid Blocker Super PAP Pen (Daido Sangyo) was used to circle two parts of a microscopy slide (VWR, 631-1161) and membranes were cut out form the inserts, using a scalpel, and placed within the PAP circle. Overnight primary antibody incubation at 4 °C was performed, in a humidified chamber. After incubation, membranes were washed three times with Tris-buffered saline (TBS)-Tween (150 mM NaCl, 50 mM Tris-HCL buffer and 1% w/v Tween20) (TBS; EMD Millipore 524750, Tween20; Sigma-Aldrich) followed by secondary antibody and Hoechst 33342 (Invitrogen, H3570, 1:1000 in Fish Serum Blocking Buffer) incubation for 2 h at RT. After three washes with PBS, membranes were mounted using ProLong TM Glass Antifade Mountant (ThermoFisher Scientific, P36984). A EVOS TM M5000 or Leica TCS SP8-X microscope with HC Plan Apochromat 40x and 63x oil objective were used to image the slides. Leica LAS X Software (Leica Microsystems) was used for analysis.

Human adult intestinal mucosa cultures were fixed in 4 % (v/v) paraformaldehyde (PFA) or cold methanol for 30 and 10 min respectively at RT and stored in PBS at 4 °C. According to the target localisation, cells were permeabilised with 0.2 % Triton-X 100 in PBS for 10 min at room temperature. Treatment with 1% BSA in PBS for 60 min was used to block non-specific binding. Subsequently, cultures were treated with primary antibodies at specific dilutions for each target (ranging from 1:50 to 1:400) for 60 min at RT. Cultures were then incubated with secondary antibodies (dilution 1:400) in the dark for 60 min and counterstained with 4′,6- diamidino-2-phenylindole (DAPI) for 1 min. Lastly, cultures were left with 0.5 mL PBS for observation with Leica DMi8 inverted fluorescence microscope (Leica Microsystems, Germany). Mean fluorescent intensity per image was quantified using ImageJ software (NIH, USA), and normalised to the number of cells per image indicated by DAPI stain. A list of the antibodies used in this study can be found in **Supplementary Table 2**.

### 5.12. Flow cytometry analysis

Cells were enzymatically dissociated with TrypLE Express for 15 min at RT. The collected cell suspension was then mechanically dissociated followed by the addition of 2 mL FACS buffer (PBS supplemented with 0.1% BSA) was added. Cells were centrifuged at 300 x g for 5 min at 4°C, and the supernatant was discarded. The pellet was resuspended in 1 mL FACS buffer, and the cells counted with an automatic cell counter. For each flow cytometry assay, 250,000 live cells were distributed in each sample tube. If epitope localisation was intracellular, cells were fixed in 4 % PFA or cold methanol for 10 min at RT, followed by permeabilised with 0.2 % Triton-X 100 in PBS for 10 min at RT. Blocking was performed using Human TruStain FcX™, Fc Receptor Blocking Solution for 10 min at RT. Then, cells were stained with conjugated antibody cocktails diluted in 100 µL FACS buffer for 45 min in the dark at RT. Unless otherwise stated, DAPI was used as a viability stain for 1 min at RT. Following staining samples were washed three times in 2 mL FACS buffer followed by centrifugation at 300 x g for 5 min. Samples were then resuspended in 100 µL FACS buffer and added to a 96-well plate. At least 10,000 cells were analysed using CytoFlex (Beckman Coulter, USA) together with CytExpert Software (Beckman Coulter, USA). Compensation matrices were generated for each multi- colour panel of antibodies, with the following gating strategy. The stability of each run was assessed by looking at the forward scatter area (FSC-A) in time. Eventual irregularities in the flow were gated out from the analysis. A doublet exclusion gate was also set on an FSC-A against forward scatter height (FSC-H) plot. Dead cells were eliminated from the analysis by including a live/dead stain in each assay. The remaining events represented the population of interest. A list of the conjugated antibodies used in this study can be found in **Supplementray Table 3**.

### 5.13. Statistical analysis

Cells derived from different donors were treated as biological replicates (n). Consequently, co-cultures involving IECs, IFs, and APCs sourced from different donors, and combined in all possible permutations, were also considered distinct biological replicates. In contrast, replicate wells containing cells derived from the same donor were treated as technical replicates, reflecting experimental repeats rather than biological variability. Unless otherwise specified, all experiments were conducted in triplicates, and data are expressed as mean ± standard deviation. Statistical analysis was conducted using GraphPad Prism 8 software (GraphPad Software Inc., USA). Normality test (D’Agostino-Pearson normality test) and equality of variances tests (Bonett’s test and Levene’s test) were conducted. When the assumptions of parametric analysis were confirmed, a student’s t-test was used to compare two groups, and a one-way analysis of variance (ANOVA) was used to compare more than 2 groups, followed by Bonferroni’s post-hoc test. Where more than one factor influenced the variable being measured, two-way ANOVA and Tukey’s multiple comparison tests were used to assess for a significant effect of each factor, as well as an interaction between factors. When the assumptions of parametric analysis were violated, a Mann-Whitney U test or a Kruskal-Wallis test were used to compare two or more independent groups, respectively. Statistical significance levels were denoted by asterisks, adhering to the New England Journal of Medicine (NEJM) style [p > 0.05 = (ns); p < 0.05 (*); p < 0.01 (**); p < 0.001 (***)].

## Acknowledgements

This work is co-funded by the European Union’s Horizon 2020 Research and innovation program GUTVIBRATIONS (grant 953201), the PPP allowance (Focus-on-Virus) made available by the Health Holland, Top Sector Life Sciences & Health, to the Amsterdam UMC, location Academic medical Center to stimulate public-private partnerships, and the AVATAR research grant funded by the Amsterdam Reproduction and Development institute. The funders had no role in the design of the study, data analysis, writing of the manuscript, or in the decision to publish the results. The authors gratefully acknowledge the Cambridge Advanced Imaging Centre for their support and assistance in this work. The HIS mouse facility (Amsterdam UMC, The Netherlands) is acknowledged for providing fetal tissue. The cellular imaging core facility of the Amsterdam UMC, The Netherlands is acknowledged for the advanced light microscopy support.

## Author contributions

Conceptualization, D.P., K.C.W., A.S., and S.S.; Formal Analysis, A.D., and C.C.; Funding acquisition, D.P., K.C.W, A.S., S.S.; Investigation, A.D., C.C., J.K., N.J., and E.F.; Methodology, A.D., C.C., A.S., and S.S.; Project administration, R.K.C, W.C., D.P., and K.C.W., Resources, A.E., S.L., D.P., and K.C.W.; Supervision. R.K.C., W.C., D.P., K.C.W., A.S., and S.S.; Visualization, A.D., and C.C.; Writing – original draft, A.D., C.C., A.S. and S.S.; Writing – review & editing, J.K., N.J., E.F., A.E., S.L., R.K.C., W.C., D.P., and K.C.W.

## 1. Supplementary Information

### 1.1. Supplementary Tables

**Supplementary Table 1:**
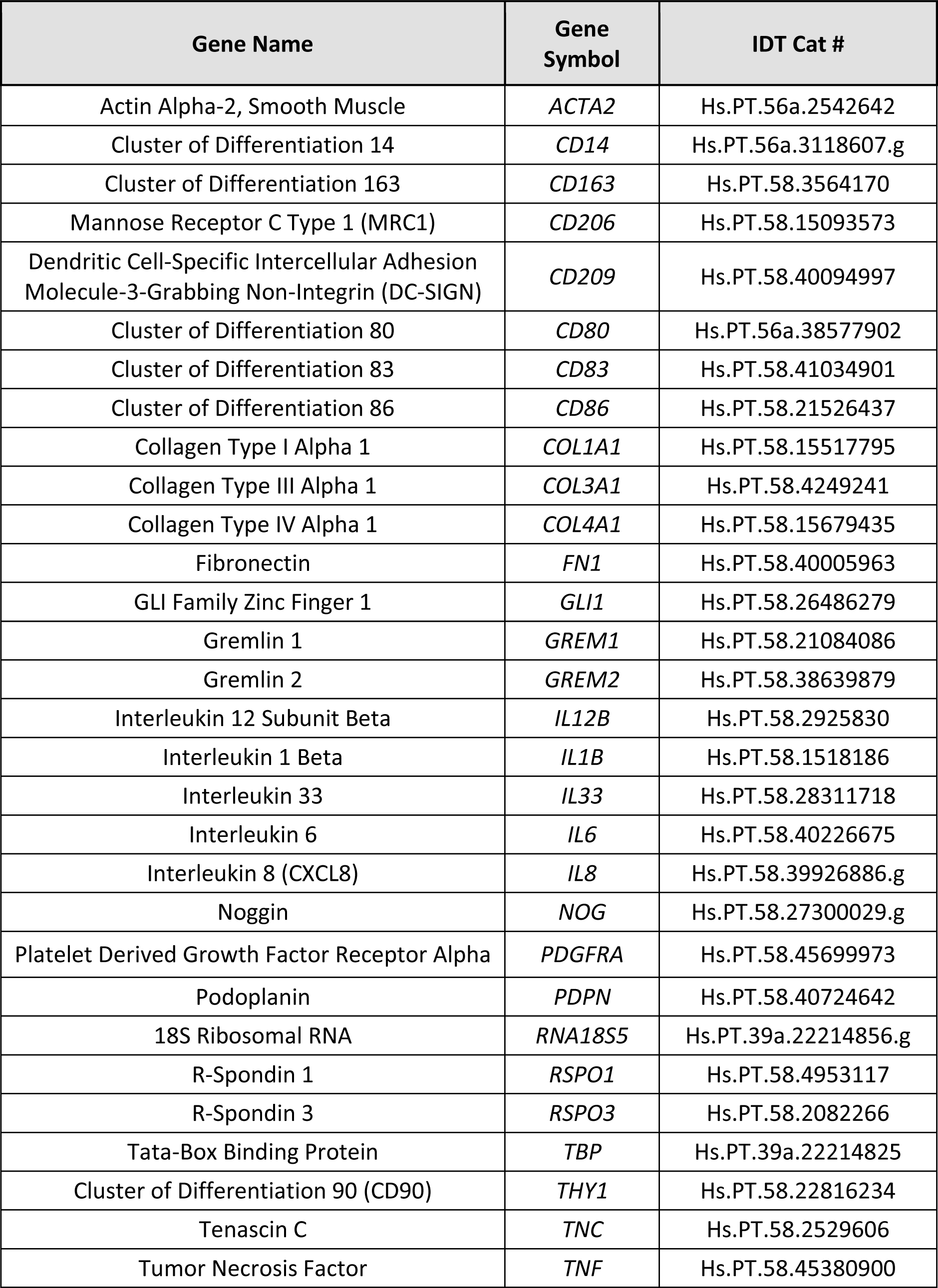
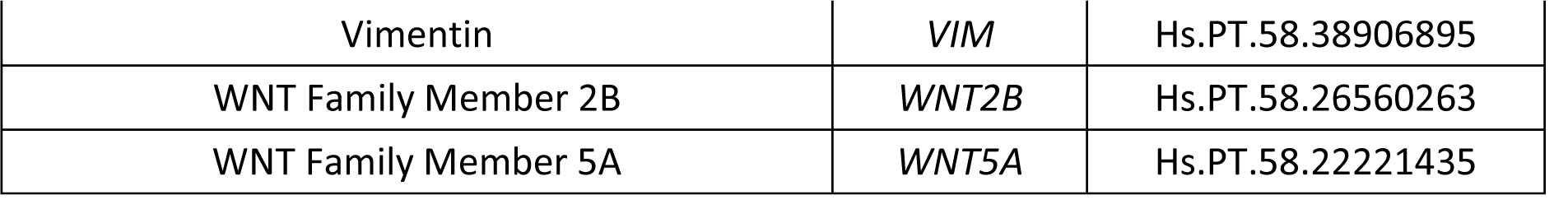
List of qPCR predesigned assays used in gene expression analysis.

| Gene Name | Gene Symbol | IDT Cat # |
| --- | --- | --- |
| Actin Alpha-2, Smooth Muscle | *ACTA2* | Hs.PT.56a.2542642 |
| Cluster of Differentiation 14 | *CD14* | Hs.PT.56a.3118607.g |
| Cluster of Differentiation 163 | *CD163* | Hs.PT.58.3564170 |
| Mannose Receptor C Type 1 (MRC1) | *CD206* | Hs.PT.58.15093573 |
| Dendritic Cell-Specific Intercellular Adhesion Molecule-3-Grabbing Non-Integrin (DC-SIGN) | *CD209* | Hs.PT.58.40094997 |
| Cluster of Differentiation 80 | *CD80* | Hs.PT.56a.38577902 |
| Cluster of Differentiation 83 | *CD83* | Hs.PT.58.41034901 |
| Cluster of Differentiation 86 | *CD86* | Hs.PT.58.21526437 |
| Collagen Type I Alpha 1 | *COL1A1* | Hs.PT.58.15517795 |
| Collagen Type III Alpha 1 | *COL3A1* | Hs.PT.58.4249241 |
| Collagen Type IV Alpha 1 | *COL4A1* | Hs.PT.58.15679435 |
| Fibronectin | *FN1* | Hs.PT.58.40005963 |
| GLI Family Zinc Finger 1 | *GLI1* | Hs.PT.58.26486279 |
| Gremlin 1 | *GREM1* | Hs.PT.58.21084086 |
| Gremlin 2 | *GREM2* | Hs.PT.58.38639879 |
| Interleukin 12 Subunit Beta | *IL12B* | Hs.PT.58.2925830 |
| Interleukin 1 Beta | *IL1B* | Hs.PT.58.1518186 |
| Interleukin 33 | *IL33* | Hs.PT.58.28311718 |
| Interleukin 6 | *IL6* | Hs.PT.58.40226675 |
| Interleukin 8 (CXCL8) | *IL8* | Hs.PT.58.39926886.g |
| Noggin | *NOG* | Hs.PT.58.27300029.g |
| Platelet Derived Growth Factor Receptor Alpha | *PDGFRA* | Hs.PT.58.45699973 |
| Podoplanin | *PDPN* | Hs.PT.58.40724642 |
| 18S Ribosomal RNA | *RNA18S5* | Hs.PT.39a.22214856.g |
| R-Spondin 1 | *RSPO1* | Hs.PT.58.4953117 |
| R-Spondin 3 | *RSPO3* | Hs.PT.58.2082266 |
| Tata-Box Binding Protein | *TBP* | Hs.PT.39a.22214825 |
| Cluster of Differentiation 90 (CD90) | *THY1* | Hs.PT.58.22816234 |
| Tenascin C | *TNC* | Hs.PT.58.2529606 |
| Tumor Necrosis Factor | *TNF* | Hs.PT.58.45380900 |
| Vimentin | *VIM* | Hs.PT.58.38906895 |
| WNT Family Member 2B | *WNT2B* | Hs.PT.58.26560263 |
| WNT Family Member 5A | *WNT5A* | Hs.PT.58.22221435 |

**Supplementary Table 2:** List of antibodies used for immunocytochemistry analysis.

| Description | Cat# | Supplier |
| --- | --- | --- |
| Anti-Collagen I antibody | ab90395 | Abcam |
| Anti-Collagen III antibody | ab7778 | Abcam |
| Anti-Collagen IV antibody | ab6586 | Abcam |
| Anti-Fibronectin antibody | ab2413 | Abcam |
| Anti-PDGFR alpha antibody [EPR22059-270] | ab203491 | Abcam |
| Anti-RSPO3 antibody | ab233113 | Abcam |
| Anti-α smooth muscle actin antibody | ab5694 | Abcam |
| APC anti-human CD103 | 350216 | Biolegend |
| EPCAM | AF960 | R&D Systems |
| FITC anti-human CD90 | 328107 | Biolegend |
| GLI1 | MA5-38530 | Invitrogen |
| Goat anti-Mouse IgG1, Alexa Fluor 647 | A-21240 | Invitrogen |
| Goat anti-Rabbit IgG, Alexa Fluor 488 | A-11034 | Invitrogen |
| IE1 |  | Gifted by Prof. Dr. William Britt, The University of Alabama |
| Lysozyme EC 3.2.1.17 | A0099 | Dako |
| PE anti-human CD108 | 376604 | Biolegend |
| Recombinant Anti-Podoplanin / gp36 antibody [EPR7072] | ab128994 | Abcam |
| RSPO1 | PA5-121183 | Invitrogen |
| Anti-Human CD45 Antibody, Clone HI30 (100 tests) | 60018AZ | STEMCELL Technologies |
| ZO-1 Monoclonal Antibody (ZO1-1A12), Alexa Fluor 488 | 339188 | Thermo Fisher Scientific |

**Supplementary Table 3:** List of antibodies used for flow cytometry analysis.

| Description | Cat# | Supplier |
| --- | --- | --- |
| Anti-DC-SIGN antibody [UW60.1] (APC) | ab180548 | Abcam |
| Anti-Human CD14 Antibody, Clone M5E2 | 60004PE | STEMCELL Technologies |
| APC anti-human CD103 | 350216 | Biolegend |
| APC anti-human CD83 | 305312 | Biolegend |
| FITC anti-human CD86 | 374204 | Biolegend |
| PE anti-human CD108 | 376604 | Biolegend |
| PE anti-human CD45 | 304007 | Biolegend |
| PE anti-human CD80 | 375410 | Biolegend |
| Recombinant Anti-Mannose Receptor antibody [EPR6828(B)] (Alexa Fluor® 488) | ab195191 | Abcam |
| Anti-Human CD45 Antibody, Clone HI30 (100 tests) | 60018AZ | STEMCELL Technologies |

### 1.2. Supplementary Figures

**Supplementary Figure 1:**
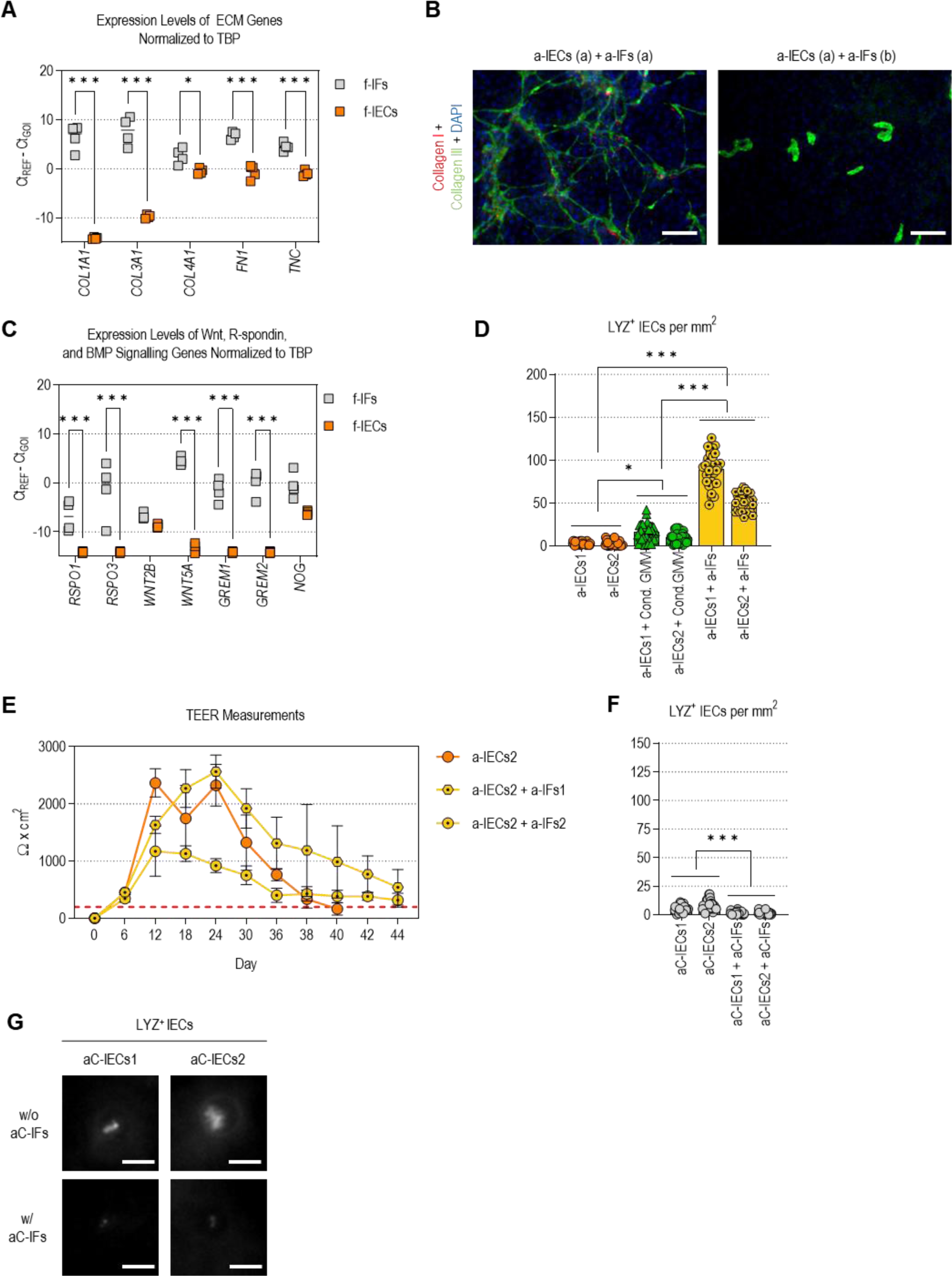
Additional data on the epithelial-mesenchymal crosstalk. (A) RT-qPCR analysis of ECM genes performed on f-IECs and f-IF monocultures maintained for 14 days on cell culture inserts in GMM. Gene expression levels are plotted as -ΔCt values. Each point represents a biological replicate (f-IFs donors: n = 4; f- IECs donors: n = 4), grey and orange colours highlight data related to IFs and IECs, respectively. Gene names are listed at the bottom of the plot. Statistical analysis employed a two-way ANOVA, along with Šídák’s multiple comparison test. **(B)** Representative immunofluorescence images of collagen type I and collagen type III expression at day 14 in IEC-IF co-cultures (2 a-IECs donors combined with 2 a-IFs donors in all possible permutations, n = 4) seeded either together on the same side [IECs (a) + IFs (a)] or on opposite sides of the insert membrane [IECs (a) + IFs (b)] (scale bars = 100 µm). **(C)** RT-qPCR analysis of WNT, R-Spondin, and BMP signalling genes performed on f-IECs and f-IFs cultures maintained for 14 days on cell culture inserts in the same medium. Gene expression levels are plotted as -ΔCt values. Each point represents a biological replicate (f-IFs donors: n = 4; f-IECs donors: n = 4), grey and orange colours highlight data related to IFs and IECs, respectively. Gene names are listed at the bottom of the plot. Statistical analysis employed a two-way ANOVA, along with Šídák’s multiple comparison test. **(D)** Bar graph summarising the results of the Paneth cells quantification based on LYZ immunostaining. The bars represent the mean and standard deviation. For each sample, at least 30 images at 10X magnification per insert were analysed. Statistical analysis employed a one-way ANOVA, along with Tukey’s multiple comparison test. **(E)** TEER measurements performed on additional IECs (n = 1), and IEC-IF co-cultures (1 a-IECs donor combined with 2 a-IFs donors, n = 2) maintained in GMM. TEER measurements were normalised to the surface area of the cell culture inserts (0.332 cm^2^) and plotted as Ω·cm^2^ (y-axis) over time (x-axis). Each point represents the mean of at least two measurements performed on different inserts, and the bars indicate the standard deviation of these measurements. Orange, and yellow colours represent data pertaining to IECs and IEC-IF co-cultures. Dotted circle and hexagon symbols denote data related to IEC-IF co-cultures incorporating IFs sourced from different donors. **(F)** Bar graph summarising the results of the Paneth cells quantification on adult colon IECs (aC-IECs) and adult colon IEC+IF co-cultures (aC-IECs + aC-IFs), based on LYZ immunostaining. The bars represent the mean and standard deviation. For each sample, at least 30 images at 10X magnification per insert were analysed. Statistical analysis employed a one-way ANOVA, along with Tukey’s multiple comparison test. **(G)** Representative immunofluorescence images of LYZ expression at day 14 in aC-IECs, and aC- IEC + aC-IF co-cultures maintained in GMM (scale bars = 10 µm).

**Supplementary Figure 2.**
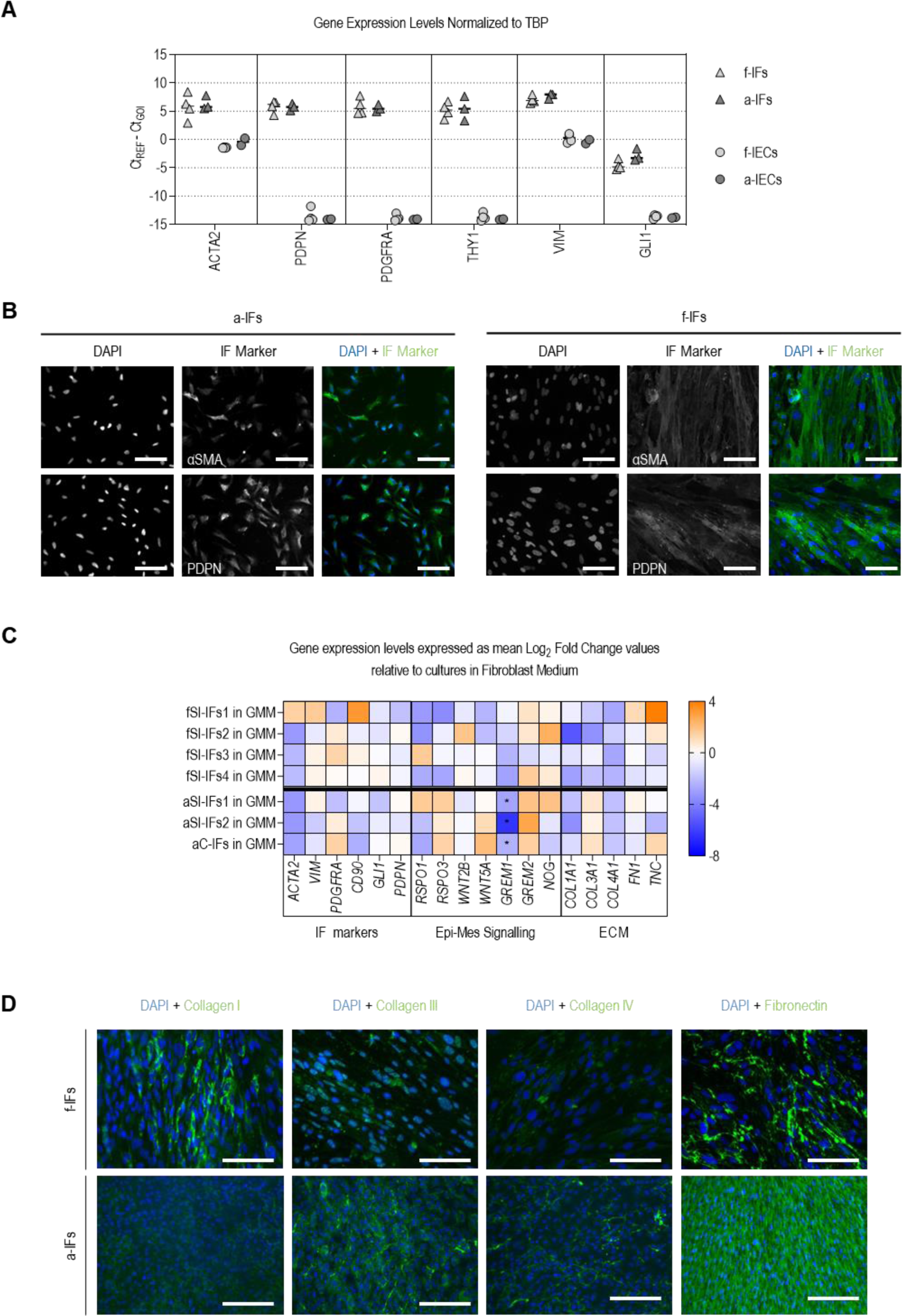
IFs expressed markers characteristic of their identity, and maintained their phenotypical and functional features in GMM. (A) Gene expression analysis by RT-qPCR on IFs cultured for 14 days in cell culture inserts in GMM. Gene expression levels are plotted as -ΔCt values. Each point represents a biological replicate (f-IFs donors: n = 4; a-IFs donors: n = 3; f-IECs donors: n = 4, a-IECs donors: n = 2), triangle and circle symbols indicate data relative to IFs and IECs respectively, while light and dark grey colours highlight data related to foetal and adult cells respectively. Gene names are listed at the bottom of the plot. **(B)** Representative immunofluorescence images of αSMA and PDPN expression in a-IFs (n = 3) and f-IFs (n = 4) cultured for 14 days in cell culture inserts in GMM (scale bars = 100 µm). For both adult and foetal IFs, the panel of 3 images shows from left to right: (i) DAPI staining the cell nuclei; (ii) the protein of interest; and (iii) the overlay of these 2 channels with blue and green highlighting the DAPI and target protein, respectively. **(C)** Heatmap representing the Log2 fold-change (Log2FC) values in gene expression of a-IF (n = 3) and f-IF (n = 4) cultures maintained for 14 days on cell culture inserts in GMM, compared to their standard culture conditions in fibroblast medium. Orange colour indicates upregulation in gene expression (Log2 fold change > 0), blue colour indicates downregulation (Log2 fold change < 0). Gene names are listed at the bottom of the heat map and are clustered by categories, related to IFs markers, epithelial-mesenchymal cell signalling, and ECM components. **(D)** Representative immunofluorescence images of collagen type I, collagen type III, collagen type IV, and fibronectin expression in a-IF (n = 3) and f-IF (n = 4) cultures maintained for 14 days on cell culture inserts in GMM (scale bars = 100 µm).

**Supplementary Figure 3.**
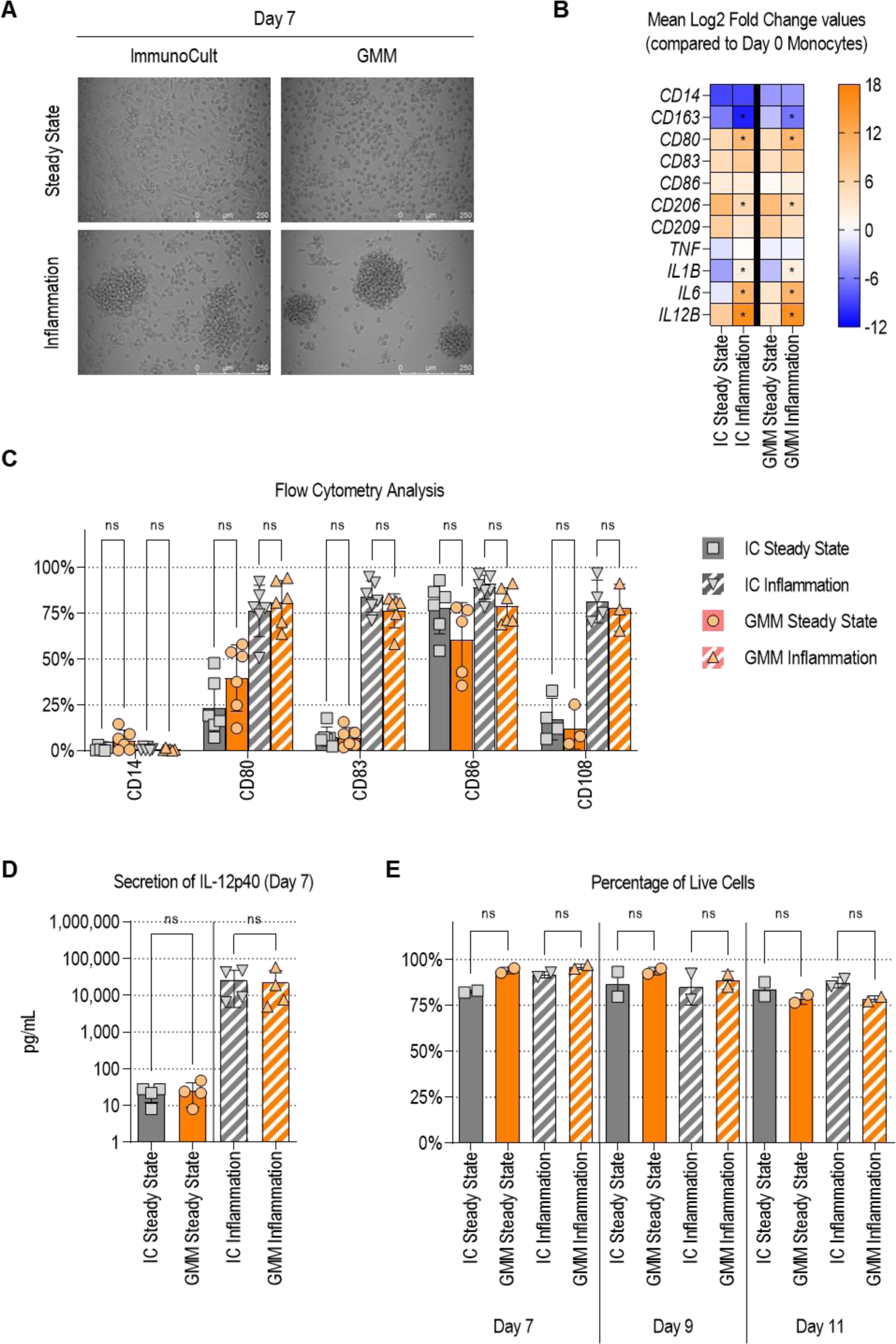
DCs retain phenotype and functions in GMM. (A) Representative phase contrast images of DCs cultured in ImmunoCult™ DC Differentiation Medium (IC) or gut mucosa medium (GMM) (n = 6; scale bars = 250 µm). **(B)** Analysis of gene expression by qPCR in DCs at day 7 cultured in different media formulations. All samples were compared to monocyte progenitors to calculate the relative gene expression (mean ± SD, n = 4) and expressed as Log2 fold changes. Orange colour indicates upregulation in gene expression (Log2 fold change > 0), blue colour indicates downregulation (Log2 fold change < 0). **(C)** Flow cytometry analysis of CD14, CD80, CD83, CD86, and CD108 protein levels at day 7 in DCs cultured in either IC or GMM, comparing steady state and inflammation conditions. Each dot represents a biological replicate (n = 6). The bars represent the mean and the standard deviation. **(D)** Dot plot indicating the concentration of IL-12p40 secreted by DCs cultured in either IC or GMM, comparing steady state and inflammation conditions. Each dot indicates a biological replicate (n = 4) and the bars represent the mean and the standard deviation. **(E)** Cell viability assessed at day 7, 9 and 11 by AO-DAPI stain in DCs cultured in either IC or GMM, comparing steady state and inflammation conditions. Each dot represents a biological replicate (n = 2) and the bars represent the mean and the standard deviation. Media conditions analysed are listed below.

**Supplementary Figure 4.**
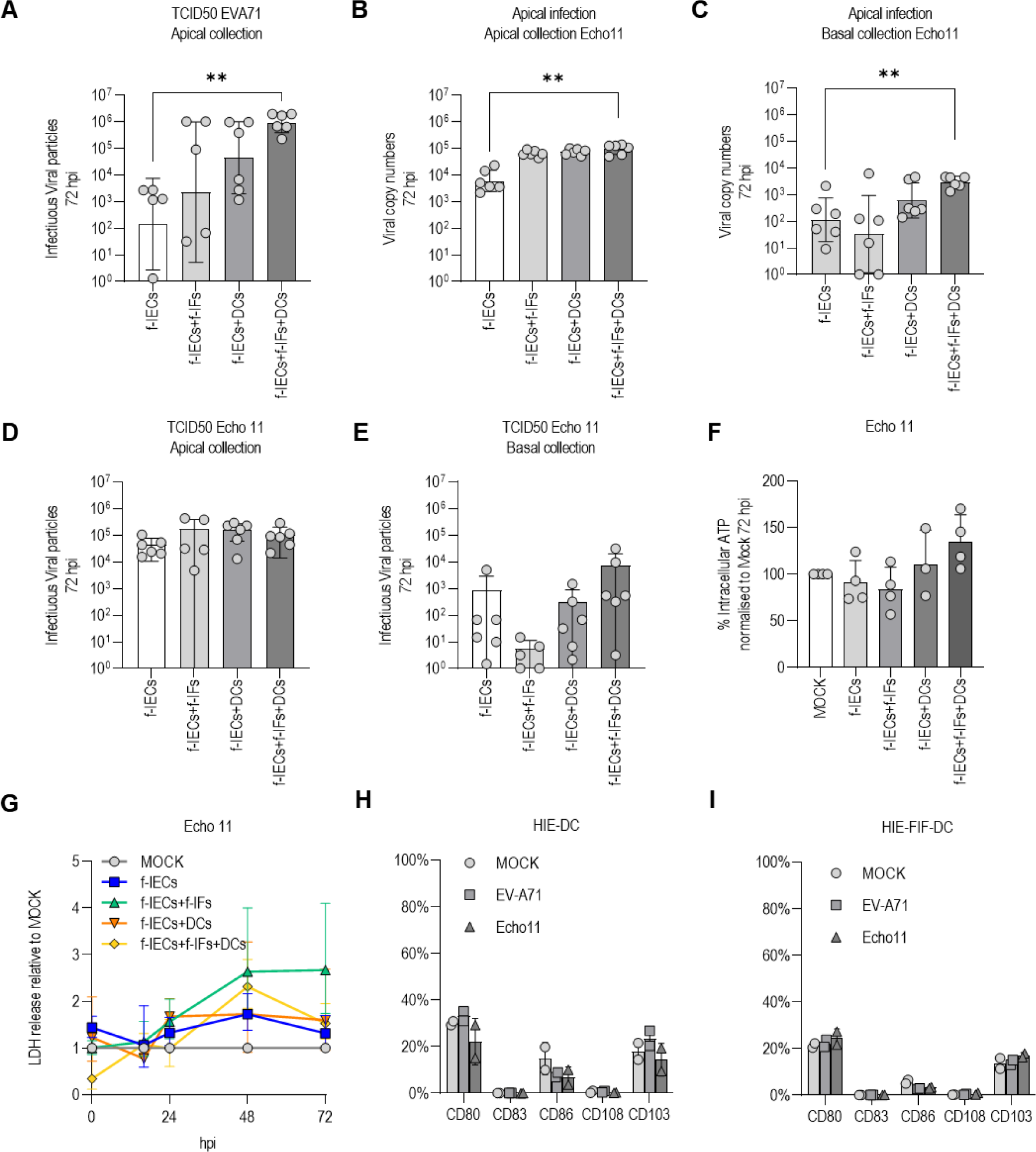
EV-A71 and Echo 11 infection in a foetal monocultures, co-cultures and gut mucosa model. (A) Average infectious viral particles in apical collected medium following apical inoculation of EV-A71, 72 hours post-infection. Data points on the graph represent results from three independent experiments across 2 biological replicates (n = 2), with bars representing the mean and standard deviation. **(B)** Average viral copy numbers in apical-collected medium following apical inoculation of Echo11, 72 hours post-infection. Data points on the graph represent results from three independent experiments across 2 biological replicates (n = 2), with bars representing the mean and standard deviation. **(C)** Average viral copy numbers in basal-collected medium following apical inoculation of Echo11, 72 hours post-infection. Data points on the graph represent results from three independent experiments across 2 biological replicates (n = 2), with bars representing the mean and standard deviation. **(D)** Average infectious viral particles in apical collected medium following apical inoculation of Echo11, 72 hours post-infection. Data points on the graph represent results from three independent experiments across 2 biological replicates (n = 2), with bars representing the mean and standard deviation. **(E)** Average infectious viral particles in basal-collected medium following apical inoculation of Echo11, 72 hours post-infection. Data points on the graph represent results from three independent experiments across 2 biological replicates (n = 2), with bars representing the mean and standard deviation. **(F)** Average increase (%) of intracellular ATP indicative of metabolic activity. Data points on the graph represent results from two independent experiments across 2 biological replicates (n = 2), with bars representing the mean and standard deviation. **(G)** LDH release relative to the uninfected mock condition for all models over time. Data points on the graph represent results from two independent experiments across 2 biological replicates (n = 2), with error bars representing the standard deviation. **(H)** Flow cytometry analysis of CD80, CD83, CD86, CD108, and CD103 expression levels following 72h EV-A71 and Echo 11 infection of IEC-DC co-cultures. Each point represents a biological replicate (n = 2, error bars = SD). **(I)** Flow cytometry analysis of CD80, CD83, CD86, CD108, and CD103 expression levels following 72h EV-A71 and Echo 11 infection of IEC-IF-DC cultures. Each point represents a biological replicate (n = 2, error bars = SD). Statistical analysis employed multiple t tests along with Bonferroni- Dunn multiple comparison tests.

## References

[1] J.W. Hickey, W.R. Becker, S.A. Nevins, A. Horning, A.E. Perez, C. Zhu, B. Zhu, B. Wei, R. Chiu, D.C. Chen, D.L. Cotter, E.D. Esplin, A.K. Weimer, C. Caraccio, V. Venkataraaman, C.M. Schürch, S. Black, M. Brbić, K. Cao, S. Chen, W. Zhang, E. Monte, N.R. Zhang, Z. Ma, J. Leskovec, Z. Zhang, S. Lin, T. Longacre, S.K. Plevritis, Y. Lin, G.P. Nolan, W.J. Greenleaf, M. Snyder, Organization of the human intestine at single-cell resolution, Nature 619(7970) (2023) 572- 584.

[2] M. Antfolk, K.B. Jensen, A bioengineering perspective on modelling the intestinal epithelial physiology in vitro, Nat. Commun. 11(1) (2020) 6244.

[3] G. Zhu, J. Hu, R. Xi, The cellular niche for intestinal stem cells: a team effort, Cell Regen. 10(1) (2021) 1.

[4] W.D. Rees, L.M. Sly, T.S. Steiner, How do immune and mesenchymal cells influence the intestinal epithelial cell compartment in inflammatory bowel disease? Let’s crosstalk about it!, J. Leukoc. Biol. 108(1) (2020) 309–321.

[5] L.E. Smythies, M. Sellers, R.H. Clements, M. Mosteller-Barnum, G. Meng, W.H. Benjamin, J.M. Orenstein, P.D. Smith, Human intestinal macrophages display profound inflammatory anergy despite avid phagocytic and bacteriocidal activity, J. Clin. Invest. 115(1) (2005) 66–75.

[6] A.A. Filardy, J.R.M. Ferreira, R.M. Rezende, B.L. Kelsall, R.P. Oliveira, The intestinal microenvironment shapes macrophage and dendritic cell identity and function, Immunol. Lett. 253 (2023) 41–53.

[7] D.S. Shouval, A. Biswas, J.A. Goettel, K. McCann, E. Conaway, N.S. Redhu, I.D. Mascanfroni, Z. Al Adham, S. Lavoie, M. Ibourk, D.D. Nguyen, J.N. Samsom, J.C. Escher, R. Somech, B. Weiss, R. Beier, L.S. Conklin, C.L. Ebens, F.G. Santos, A.R. Ferreira, M. Sherlock, A.K. Bhan, W. Müller, J.R. Mora, F.J. Quintana, C. Klein, A.M. Muise, B.H. Horwitz, S.B. Snapper, Interleukin-10 receptor signaling in innate immune cells regulates mucosal immune tolerance and anti-inflammatory macrophage function, Immunity 40(5) (2014) 706–719.

[8] D.T. Ruane, E.C. Lavelle, The role of CD103⁺ dendritic cells in the intestinal mucosal immune system, Front. Immunol. 2 (2011) 1–6.

[9] S.J. Gaudino, P. Kumar, Cross-talk between antigen presenting cells and T cells impacts intestinal homeostasis, bacterial infections, and tumorigenesis, Front. Immunol. 10 (2019) 360.

[10] W. Liu, Q. Wang, Y. Bai, H. Xiao, Z. Li, Y. Wang, Q. Wang, J. Yang, H. Sun, Potential application of intestinal organoids in intestinal diseases, Stem Cell Rev. Rep. 20(1) (2024) 124–137.

[11] L. Wu, Y. Ai, R. Xie, J. Xiong, Y. Wang, Q. Liang, Organoids/organs-on-a-chip: New frontiers of intestinal pathophysiological models, Lab Chip 23(5) (2023) 1192–1212.

[12] Y. Wang, H. Lin, L. Zhao, F. Hong, J. Hao, Z. Zhang, W. Sheng, L. Song, C.-X. Deng, B. Zhao, J. Cao, L. Wang, L. Wang, L. Liang, W.K. Chen, C. Yu, Z. Sun, Y. Yang, C. Wang, Y. Zhang, Q. Li, K. Li, A. Ma, T. Zhao, G. Hua, Y.-G. Chen, Standard: Human intestinal organoids, Cell Regen. 12(1) (2023) 1–5.

[13] W.Y. Zou, S.E. Blutt, S.E. Crawford, K. Ettayebi, X.-L. Zeng, K. Saxena, S. Ramani, U.C. Karandikar, N.C. Zachos, M.K. Estes, Human intestinal enteroids: New models to study gastrointestinal virus infections, in: K. Turksen (Ed.), Organoids: Stem Cells, Structure, and Function, Springer New York, New York, NY, 2019, pp. 229-247.

[14] J. Taelman, M. Diaz, J. Guiu, Human Intestinal Organoids: Promise and Challenge, Front. Cell Dev. Biol. 10 (2022) 1–9.

[15] T. Roodsant, M. Navis, I. Aknouch, I.B. Renes, R.M. van Elburg, D. Pajkrt, K.C. Wolthers, C. Schultsz, K.C.H. van der Ark, A. Sridhar, V. Muncan, A human 2D primary organoid-derived epithelial monolayer model to study host-pathogen interaction in the small intestine, Front. Cell. Infect. Microbiol. 10 (2020) 1–14.

[16] G. Stroulios, M. Stahl, F. Elstone, W. Chang, S. Louis, A. Eaves, S. Simmini, R.K. Conder, Culture methods to study apical-specific interactions using intestinal organoid models, JoVE (169) (2021) e62330.

[17] M. Nikolaev, O. Mitrofanova, N. Broguiere, S. Geraldo, D. Dutta, Y. Tabata, B. Elci, N. Brandenberg, I. Kolotuev, N. Gjorevski, H. Clevers, M.P. Lutolf, Homeostatic mini-intestines through scaffold-guided organoid morphogenesis, Nature 585(7826) (2020) 574-578.

[18] M. Verhulsel, A. Simon, M. Bernheim-Dennery, V.R. Gannavarapu, L. Geremie, D. Ferraro, D. Krndija, L. Talini, J.L. Viovy, D.M. Vignjevic, S. Descroix, Developing an advanced gut on chip model enabling the study of epithelial cell/fibroblast interactions, Lab Chip 21(2) (2021) 365–377.

[19] G. Weindl, Immunocompetent human intestinal models in preclinical drug development, in: M. Schäfer-Korting, S. Stuchi Maria-Engler, R. Landsiedel (Eds.), Organotypic Models in Drug Development, Springer International Publishing, Cham, 2021, pp. 219-233.

[20] A.A. Akhtar, S. Sances, R. Barrett, J.J. Breunig, Organoid and organ-on-a-chip systems: New paradigms for modeling neurological and gastrointestinal disease, Curr. Stem Cell Rep. 3(2) (2017) 98–111.

[21] M.F. Graham, R.F. Diegelmann, C.O. Elson, W.J. Lindblad, N. Gotschalk, S. Gay, R. Gay, Collagen content and types in the intestinal strictures of Crohn’s disease, Gastroenterology 94(2) (1988) 257–265.

[22] M.D. Brügger, K. Basler, The diverse nature of intestinal fibroblasts in development, homeostasis, and disease, Trends Cell Biol. 33(10) (2023) 834–849.

[23] X. Mei, M. Gu, M. Li, Plasticity of Paneth cells and their ability to regulate intestinal stem cells, Stem Cell Res. Ther. 11(1) (2020) 349.

[24] S. Yu, I. Balasubramanian, D. Laubitz, K. Tong, S. Bandyopadhyay, X. Lin, J. Flores, R. Singh, Y. Liu, C. Macazana, Y. Zhao, F. Béguet-Crespel, K. Patil, M.T. Midura-Kiela, D. Wang, G.S. Yap, R.P. Ferraris, Z. Wei, E.M. Bonder, M.M. Häggblom, L. Zhang, V. Douard, M.P. Verzi, K. Cadwell, P.R. Kiela, N. Gao, Paneth cell-derived lysozyme defines the composition of mucolytic microbiota and the inflammatory tone of the intestine, Immunity 53(2) (2020) 398–416.

[25] C.A. Thomson, R.J. Nibbs, K.D. McCoy, A.M. Mowat, Immunological roles of intestinal mesenchymal cells, Immunology 160(4) (2020) 313–324.

[26] A. Parker, L. Vaux, A.M. Patterson, A. Modasia, D. Muraro, A.G. Fletcher, H.M. Byrne, P.K. Maini, A.J.M. Watson, C. Pin, Elevated apoptosis impairs epithelial cell turnover and shortens villi in TNF-driven intestinal inflammation, Cell Death Discov. 10(2) (2019) 108.

[27] C. Sinzger, A. Grefte, B. Plachter, A.S. Gouw, T.H. The, G. Jahn, Fibroblasts, epithelial cells, endothelial cells and smooth muscle cells are major targets of human cytomegalovirus infection in lung and gastrointestinal tissues, J. Gen. Virol. 76 ( Pt 4) (1995) 741–750.

[28] P.-J. Yeh, R.-C. Wu, C.-L. Chen, C.-T. Chiu, M.-W. Lai, C.-C. Chen, C.-H. Chiu, Y.-B. Pan, W.- R. Lin, P.-H. Le, Cytomegalovirus diseases of the gastrointestinal tract in immunocompetent patients: A narrative review, Viruses, 2024, pp. 1–17.

[29] V.T.K. Le-Trilling, J.F. Ebel, F. Baier, K. Wohlgemuth, K.R. Pfeifer, A. Mookhoek, P. Krebs, M. Determann, B. Katschinski, A. Adamczyk, E. Lange, R. Klopfleisch, C.M. Lange, V. Sokolova, M. Trilling, A.M. Westendorf, Acute cytomegalovirus infection modulates the intestinal microbiota and targets intestinal epithelial cells, Eur. J. Immunol. 53(2) (2023) e2249940.

[30] Y.-D. Xu, M. Cheng, P.-P. Shang, Y.-Q. Yang, Role of IL-6 in dendritic cell functions, J. Leukoc. Biol. 111(3) (2022) 695–709.

[31] R. Marone, S. Garaudé, R. Lepore, A. Devaux, A. Beerlage, F. Simonetta, A. Camus, I. Durzynska, I. Kirby, P. Van Berkel, C. Kunz, S. Urlinger, L.T. Jeker, Hematopoietic stem cells expressing engineered CD45 enable a near universal targeted therapy for hematologic diseases, Blood 142(Supplement 1) (2023) 3418–3418.

[32] C.L. Yao, T.Y. Tseng, The synergistic and enhancive effects of IL-6 and M-CSF to expand and differentiate functional dendritic cells from human monocytes under serum-free condition, J. Biol. Eng. 17(1) (2023) 6.

[33] C. Koorella, J.R. Nair, M.E. Murray, L.M. Carlson, S.K. Watkins, K.P. Lee, Novel regulation of CD80/CD86-induced phosphatidylinositol 3-kinase signaling by NOTCH1 protein in interleukin-6 and indoleamine 2,3-dioxygenase production by dendritic cells, J. Biol. Chem. 289(11) (2014) 7747–7762.

[34] J.M. Bates, K. Flanagan, L. Mo, N. Ota, J. Ding, S. Ho, S. Liu, M. Roose-Girma, S. Warming, L. Diehl, Dendritic cell CD83 homotypic interactions regulate inflammation and promote mucosal homeostasis, Mucosal Immunol. 8(2) (2015) 414–428.

[35] T.S. Lim, J.K. Goh, A. Mortellaro, C.T. Lim, G.J. Hämmerling, P. Ricciardi-Castagnoli, CD80 and CD86 differentially regulate mechanical interactions of T-cells with antigen-presenting dendritic cells and B-cells, PLoS One 7(9) (2012) e45185.

[36] Z. Li, X. Ju, P.A. Silveira, E. Abadir, W.-H. Hsu, D.N.J. Hart, G.J. Clark, CD83: Activation marker for antigen presenting cells and its therapeutic potential, Front. Immunol. 10 (2019) 1–9.

[37] A. van Rijn, L. Paulis, J. te Riet, A. Vasaturo, I. Reinieren-Beeren, A. van der Schaaf, A.J. Kuipers, L.P. Schulte, B.C. Jongbloets, R.J. Pasterkamp, C.G. Figdor, A.B. van Spriel, S.I. Buschow, Semaphorin 7A promotes chemokine-driven dendritic cell migration, J. Immunol. 196(1) (2016) 459–468.

[38] O. Annacker, J.L. Coombes, V. Malmstrom, H.H. Uhlig, T. Bourne, B. Johansson-Lindbom, W.W. Agace, C.M. Parker, F. Powrie, Essential role for CD103 in the T cell-mediated regulation of experimental colitis, J. Exp. Med. 202(8) (2005) 1051–1061.

[39] I. Aknouch, I. García-Rodríguez, F.P. Giugliano, C. Calitz, G. Koen, H. van Eijk, N. Johannessson, S. Rebers, L. Brouwer, V. Muncan, K.J. Stittelaar, D. Pajkrt, K.C. Wolthers, A. Sridhar, Amino acid variation at VP1-145 of enterovirus A71 determines the viral infectivity and receptor usage in a primary human intestinal model, Front. Microbiol. 14 (2023) 1–10.

[40] L.C. Helgers, M.S. Bhoekhan, D. Pajkrt, K.C. Wolthers, T.B.H. Geijtenbeek, A. Sridhar, Human dendritic cells transmit Enterovirus A71 via heparan sulfates to target cells independent of viral replication, Microbiol. Spectr. 10(6) (2022) e02822–22.

[41] C. Wang, R. Yang, F. Yang, Y. Han, Y. Ren, X. Xiong, X. Wang, Y. Bi, L. Li, Y. Qiu, Y. Xu, X. Zhou, Echovirus 11 infection induces pyroptotic cell death by facilitating NLRP3 inflammasome activation, PLOS Pathogens 18(8) (2022) e1010787.

[42] L. Wu, Z. Yan, Y. Jiang, Y. Chen, J. Du, L. Guo, J. Xu, Z. Luo, Y. Liu, Metabolic regulation of dendritic cell activation and immune function during inflammation, Front. Immunol. 14 (2023) 1140749.

[43] S.H. Lee, P.M. Starkey, S. Gordon, Quantitative analysis of total macrophage content in adult mouse tissues. Immunochemical studies with monoclonal antibody F4/80, J. Exp. Med. 161(3) (1985) 475-489.

[44] Q. Wei, Y. Deng, Q. Yang, A. Zhan, L. Wang, The markers to delineate different phenotypes of macrophages related to metabolic disorders, Front. Immunol. 14 (2023) 1084636.

[45] A. Mantovani, A. Sica, S. Sozzani, P. Allavena, A. Vecchi, M. Locati, The chemokine system in diverse forms of macrophage activation and polarization, Trends Immunol. 25(12) (2004) 677–686.

[46] C.D. Mills, K. Kincaid, J.M. Alt, M.J. Heilman, A.M. Hill, M-1/M-2 macrophages and the Th1/Th2 paradigm, J. Immunol. 164(12) (2000) 6166–6173.

[47] P.J. Murray, Macrophage polarization, Annu. Rev. Physiol. 79 (2017) 541–566.

[48] Z. Strizova, I. Benesova, R. Bartolini, R. Novysedlak, E. Cecrdlova, L.K. Foley, I. Striz, M1/M2 macrophages and their overlaps - Myth or reality?, Clin Sci (Lond) 137(15) (2023) 1067–1093.

[49] A. Vilardi, S. Przyborski, C. Mobbs, A. Rufini, C. Tufarelli, Current understanding of the interplay between extracellular matrix remodelling and gut permeability in health and disease, Cell Death Discov. 10(1) (2024) 258.

[50] M. Maimets, M.T. Pedersen, J. Guiu, J. Dreier, M. Thodberg, Y. Antoku, P.J. Schweiger, L. Rib, R.B. Bressan, Y. Miao, K.C. Garcia, A. Sandelin, P. Serup, K.B. Jensen, Mesenchymal-epithelial crosstalk shapes intestinal regionalisation via Wnt and Shh signalling, Nat. Commun. 13(1) (2022) 715.

[51] R. Thibeaux, A. Dufour, P. Roux, M. Bernier, A.C. Baglin, P. Frileux, J.C. Olivo-Marin, N. Guillén, E. Labruyère, Newly visualized fibrillar collagen scaffolds dictate Entamoeba histolytica invasion route in the human colon, Cell Microbiol. 14(5) (2012) 609–621.

[52] R. Ramadan, V.M. Wouters, S.M. van Neerven, N.E. de Groot, T.M. Garcia, V. Muncan, O.D. Franklin, M. Battle, K.S. Carlson, J. Leach, O.J. Sansom, O. Boulard, M. Chamaillard, L. Vermeulen, J.P. Medema, D.J. Huels, The extracellular matrix controls stem cell specification and crypt morphology in the developing and adult mouse gut, Biol. Open. 11(12) (2022) 1–13.

[53] C.A. Duckworth, Identifying key regulators of the intestinal stem cell niche, Biochem. Soc. Trans. 49(5) (2021) 2163–2176.

[54] H.F. Farin, J.H. Van Es, H. Clevers, Redundant sources of Wnt regulate intestinal stem cells and promote formation of Paneth cells, Gastroenterology 143(6) (2012) 1518–1529.

[55] E. Maidji, M. Somsouk, J.M. Rivera, P.W. Hunt, C.A. Stoddart, Replication of CMV in the gut of HIV-infected individuals and epithelial barrier dysfunction, PLOS Pathogens 13(2) (2017) e1006202.

[56] K.J. Cavagnero, R.L. Gallo, Essential immune functions of fibroblasts in innate host defense, Front. Immunol. 13 (2022) 1058862.

[57] M. Pasztoi, C. Ohnmacht, Tissue niches formed by intestinal mesenchymal stromal cells in mucosal homeostasis and immunity, Int. J. Mol. Sci. 23(9) (2022) 1–19.

[58] V. Jeffery, A.J. Goldson, J.R. Dainty, M. Chieppa, A. Sobolewski, IL-6 signaling regulates small intestinal crypt homeostasis, J. Immunol. 199(1) (2017) 304–311.

[59] N.R. West, Coordination of immune-stroma crosstalk by IL-6 family cytokines, Front. Immunol. 10 (2019) 1–16.

[60] T.C. Barnes, M.E. Anderson, R.J. Moots, The many faces of interleukin-6: The role of IL-6 in inflammation, vasculopathy, and fibrosis in systemic sclerosis, Int. J. Rheumatol. 2011(1) (2011) 721608.

[61] L. Welz, K. Aden, Fibrosis and inflammation in inflammatory bowel disease: More than 2 sides of the same coin?, Gastroenterology 164(1) (2023) 19–21.

[62] A. Shahini, A. Shahini, Role of interleukin-6-mediated inflammation in the pathogenesis of inflammatory bowel disease: Focus on the available therapeutic approaches and gut microbiome, J. Cell Commun. Signal. 17(1) (2023) 55–74.

[63] S. Schreiber, K. Aden, J.P. Bernardes, C. Conrad, F. Tran, H. Höper, V. Volk, N. Mishra, J.I. Blase, S. Nikolaus, J. Bethge, T. Kühbacher, C. Röcken, M. Chen, I. Cottingham, N. Petri, B.B. Rasmussen, J. Lokau, L. Lenk, C. Garbers, F. Feuerhake, S. Rose-John, G.H. Waetzig, P. Rosenstiel, Therapeutic interleukin-6 trans-signaling inhibition by olamkicept (sgp130Fc) in patients with active inflammatory bowel disease, Gastroenterology 160(7) (2021) 2354–2366.

[64] H. Ito, M. Takazoe, Y. Fukuda, T. Hibi, K. Kusugami, A. Andoh, T. Matsumoto, T. Yamamura, J. Azuma, N. Nishimoto, K. Yoshizaki, T. Shimoyama, T. Kishimoto, A pilot randomized trial of a human anti-interleukin-6 receptor monoclonal antibody in active Crohn’s disease, Gastroenterology 126(4) (2004) 989–996.

[65] S. Danese, S. Vermeire, P. Hellstern, R. Panaccione, G. Rogler, G. Fraser, A. Kohn, P. Desreumaux, R.W. Leong, G.M. Comer, F. Cataldi, A. Banerjee, M.K. Maguire, C. Li, N. Rath, J. Beebe, S. Schreiber, Randomised trial and open-label extension study of an anti-interleukin-6 antibody in Crohn’s disease (ANDANTE I and II), Gut 68(1) (2019) 40–48.

[66] S.-M. Jung, S. Kim, *In vitro* models of the small intestine for studying intestinal diseases, Front. Microbiol. 12 (2022) 1–15.

[67] A.-C.R. Beitnes, M. Ráki, K.E.A. Lundin, J. Jahnsen, L.M. Sollid, F.L. Jahnsen, Density of CD163+CD11c+ dendritic cells increases and CD103+ dendritic cells decreases in the coeliac lesion, Scand. J. Immunol. 74(2) (2011) 186–194.

[68] M. Rescigno, A. Di Sabatino, Dendritic cells in intestinal homeostasis and disease, J. Clin. Invest. 119(9) (2009) 2441–2450.

[69] S.M. Dillon, L.M. Rogers, R. Howe, L.A. Hostetler, J. Buhrman, M.D. McCarter, C.C. Wilson, Human intestinal lamina propria CD1c+ dendritic cells display an activated phenotype at steady state and produce IL-23 in response to TLR7/8 stimulation, J. Immunol. 184(12) (2010) 6612–6621.

[70] H. Matsuno, H. Kayama, J. Nishimura, Y. Sekido, H. Osawa, S. Barman, T. Ogino, H. Takahashi, N. Haraguchi, T. Hata, C. Matsuda, H. Yamamoto, M. Uchino, H. Ikeuchi, Y. Doki, M. Mori, K. Takeda, T. Mizushima, CD103+ dendritic cell function is altered in the colons of patients with ulcerative colitis, Inflamm. Bowel Dis. 23(9) (2017) 1524–1534.

[71] L. Marongiu, M. Valache, F.A. Facchini, F. Granucci, How dendritic cells sense and respond to viral infections, Clin Sci (Lond) 135(19) (2021) 2217–2242.

[72] M.-C. José Luis, C.-C. Juan Francisco, G.-C. Oscar, V.-G. Paola Trinidad, R.-G. Luis Guillermo, V.-B. Jazmín Monserrat, Role of dendritic cells in pathogen infections: A current perspective, in: S. Bhawana (Ed.), Cell Interaction, IntechOpen, Rijeka, 2021, p. Ch. 8.

[73] H.I. Huang, J.Y. Lin, H.C. Chiang, P.N. Huang, Q.D. Lin, S.R. Shih, Exosomes facilitate transmission of Enterovirus A71 from human intestinal epithelial cells, J. Infect. Dis. 222(3) (2020) 456–469.

[74] P. Siciński, J. Rowiński, J.B. Warchoł, Z. Jarzabek, W. Gut, B. Szczygieł, K. Bielecki, G. Koch, Poliovirus type 1 enters the human host through intestinal M cells, Gastroenterology 98(1) (1990) 56–58.

[75] L. Ouzilou, E. Caliot, I. Pelletier, M.-C. Prévost, E. Pringault, F. Colbère-Garapin, Poliovirus transcytosis through M-like cells, J. Gen. Virol. 83(9) (2002) 2177–2182.

[76] E. Hassan, M.T. Baldridge, Norovirus encounters in the gut: multifaceted interactions and disease outcomes, Mucosal Immunol. 12(6) (2019) 1259–1267.

[77] M.B. Paul, M. Schlief, H. Daher, A. Braeuning, H. Sieg, L. Böhmert, A human Caco-2-based co-culture model of the inflamed intestinal mucosa for particle toxicity studies, In vitro models 2(1) (2023) 43–64.

[78] S. Jiang, T. Deng, H. Cheng, W. Liu, D. Shi, J. Yuan, Z. He, W. Wang, B. Chen, L. Ma, X. Zhang, P. Gong, Macrophage-organoid co-culture model for identifying treatment strategies against macrophage-related gemcitabine resistance, J. Exp. Clin. Cancer Res. 42(1) (2023) 199.

[79] Y. Yao, W. Shang, L. Bao, Z. Peng, C. Wu, Epithelial-immune cell crosstalk for intestinal barrier homeostasis, Eur. J. Immunol. 54(6) (2024) 2350631.

[80] A. Hausmann, C. Steenholdt, O.H. Nielsen, K.B. Jensen, Immune cell-derived signals governing epithelial phenotypes in homeostasis and inflammation, Trends Mol. Med. 30(3) (2024) 239–251.

[81] N.M. Nanlohy, N. Johannesson, L. Wijnands, L. Arroyo, J. de Wit, G. den Hartog, K.C. Wolthers, A. Sridhar, S. Fuentes, Exploring host-commensal-pathogen dynamics in cell line and organotypic human intestinal epithelial models, iScience 27(5) (2024) 1–15.

[82] P. Shah, J.V. Fritz, E. Glaab, M.S. Desai, K. Greenhalgh, A. Frachet, M. Niegowska, M. Estes, C. Jäger, C. Seguin-Devaux, F. Zenhausern, P. Wilmes, A microfluidics-based in vitro model of the gastrointestinal human–microbe interface, Nat. Commun. 7(1) (2016) 11535.

[83] S. Jalili-Firoozinezhad, F.S. Gazzaniga, E.L. Calamari, D.M. Camacho, C.W. Fadel, A. Bein, B. Swenor, B. Nestor, M.J. Cronce, A. Tovaglieri, O. Levy, K.E. Gregory, D.T. Breault, J.M.S. Cabral, D.L. Kasper, R. Novak, D.E. Ingber, A complex human gut microbiome cultured in an anaerobic intestine-on-a-chip, Nat. Biomed. Eng. 3(7) (2019) 520–531.

[84] I. García-Rodríguez, G. Moreni, P.E. Capendale, L. Mulder, I. Aknouch, R. Vieira de Sá, N. Johannesson, E. Freeze, H. van Eijk, G. Koen, K.C. Wolthers, D. Pajkrt, A. Sridhar, C. Calitz, Assessment of the broad-spectrum host targeting antiviral efficacy of halofuginone hydrobromide in human airway, intestinal and brain organotypic models, Antiviral Res. 222 (2024) 105798.

[85] I. Aknouch, A. Sridhar, E. Freeze, F.P. Giugliano, B.J. van Keulen, M. Romijn, C. Calitz, I. García-Rodríguez, L. Mulder, M.E. Wildenberg, V. Muncan, M.J. van Gils, J.B. van Goudoever, K.J. Stittelaar, K.C. Wolthers, D. Pajkrt, Human milk inhibits some enveloped virus infections, including SARS-CoV-2, in an intestinal model, Life Sci Alliance 5(12) (2022) 1–15.

[86] L.J. Reed, H. Muench, A simple method of estimating fifty per cent endpoints, Am. J. Epidemiol. 27(3) (1938) 493–497.

